# Microbial photoproduction of heptane

**DOI:** 10.1101/2024.07.17.603920

**Authors:** Ángel Baca-Porcel, Bertrand Legeret, Pascaline Auroy-Tarrago, Florian Veillet, Cécile Giacalone, Stephan Cuine, Poutoum Palakiyém Samire, Solène Moulin, Yonghua Li-Beisson, Fred Beisson, Damien Sorigué

## Abstract

Fatty Acid Photodecarboxylase (FAP) has emerged as a promising catalyst for the biological production of long-chain hydrocarbons. We have recently shown that purified FAP or FAP-expressing bacteria can efficiently convert octanoic acid into heptane, thus extending the potential applications of FAP to medium-chain hydrocarbons (i.e., solvent- or kerosene-type). The scarcity of natural sources of octanoic acid presents a challenge however. Here, we explore the heptane production capacity of a FAP-expressing *E. coli* strain engineered to biosynthesize octanoic acid via a specific thioesterase. Various FAPs and C8-specific thioesterases were tested. A blue-light-inducible promoter was used to avoid chemical inducers. We found that the expression of FAP fused with TrxA resulted in a 10-fold increase in heptane production. Coexpression of *Cuphea hookeriana* thioesterase and *Chlorella variabilis* FAP achieved the highest heptane titer (12.5 mg.L^-1^). Scale-up experiments in 100 mL photobioreactors allowed a constant production of heptane over two days (22 mg.L^-1^.day^-1^).

**Graphical Abstract:** 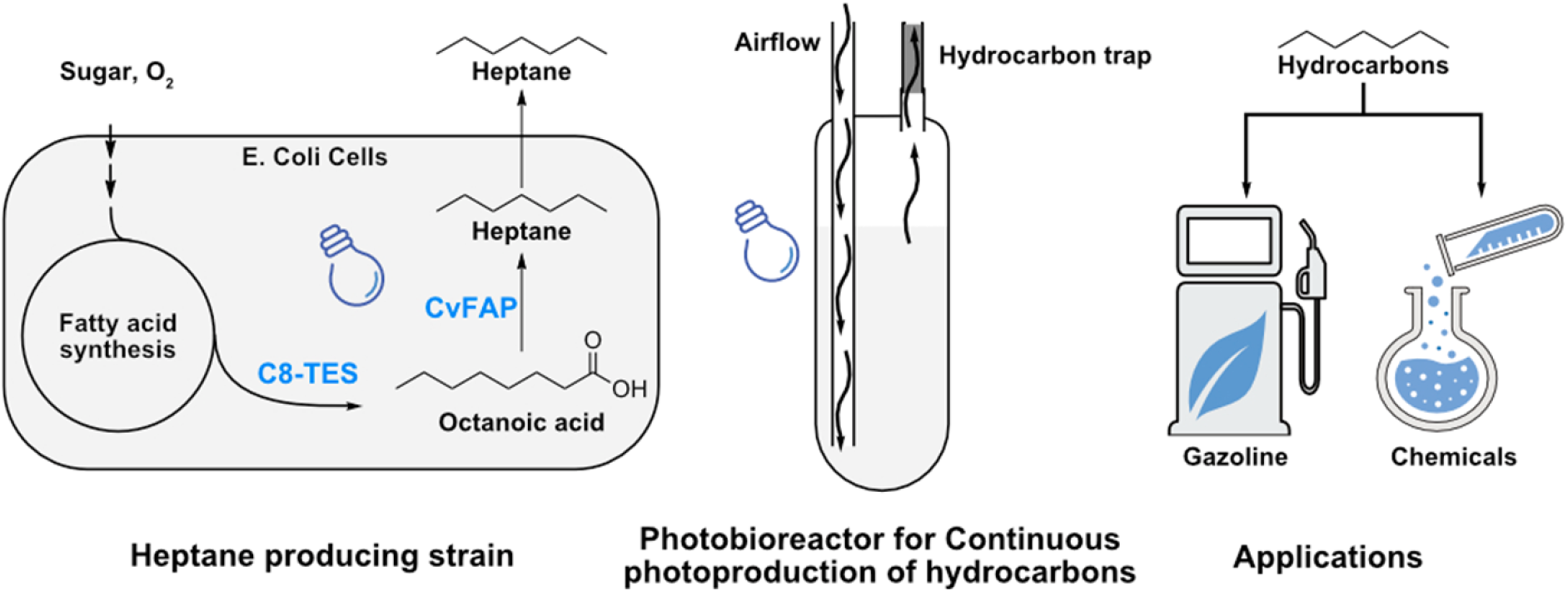

**Highlights:**

- An *E. coli* strain producing heptane under blue light is described.
- Expression of FAP fused with TrxA increases heptane by 10-fold.
- A blue light-inducible promoter ensures high coexpression of FAP and thioesterase.
- *Cuphea hookeriana* thioesterase and *Chlorella variabilis* FAP give highest production
- Highest reported heptane productivity (22 mg.L^-1^.day^-1^) in 100 mL photobioreactors.

## 1. Introduction

Beyond their function as fuel, hydrocarbons (HCs) extracted from petroleum serve as precursors to a myriad of chemicals and find applications in solvents, manufacturing and cosmetics formulations (Kuppusamy et al., n.d.; Niederer et al., 2016). Escalating global demand for fossil HCs, coupled with environmental concerns arising from their exploitation and the threat of a shortage, underlines the need to find sustainable alternative HC sources, especially for aviation and chemistry. In this context, harnessing the biological synthesis of HCs using industrial microorganisms appears to be a significant biotechnological opportunity (Geng et al., 2023).

Remarkably, nature offers a wide variety of organisms, including plants, insects, bacteria, and microalgae, that harbor enzymes capable of converting fatty acids (FAs) or their derivatives into linear alkanes or alkenes (Herman and Zhang, 2016; Jaroensuk et al., 2020). These enzymes have been expressed in yeasts, bacteria or microalgae to produce HCs (Jaroensuk et al. 2020; Geng et al. 2023). These studies (Amer et al., 2020a, 2020b; Bruder et al., 2019; Cao et al., 2016; Crépin et al., 2018, 2016; Fatma et al., 2018; Li et al., 2020; Schirmer et al., 2010; Yang et al., 2019; Yunus et al., 2022; Yunus et al., 2018), which have employed various enzymes and host strains, have been mainly focused on short-chain HCs (like propane, butane and isobutane) as well as long-chain HCs. The production of medium-chain HCs (C7-C13), which are major components in gasoline, jet fuels, and solvents (Kuppusamy et al. 2020), was also explored (Akhtar et al., 2013; Blazeck et al., 2013; Rui et al., 2015; Yan et al., 2016; Moulin et al., 2019; Yunus et al., 2022; Yunus et al., 2018) by combining the expression of an HC-forming enzyme with a specific acyl-acyl carrier protein thioesterase shortening fatty acids. However, the yield was limited and in many cases the saturated medium-chain HC is not the main product.

Among the HC-forming enzymes, the most recently discovered is fatty acid photodecarboxylase (FAP, EC 4.1.1.106), which catalyzes in algae the flavin-based photodecarboxylation of FAs into HCs within the photon wavelength range of 350 to 530 nm (Sorigué et al., 2017). The activity of this photoenzyme has been characterized *in vitro* on a wide range of fatty acids randomly dispersed in bulk or incorporated into organized lipid assemblies (Aselmeyer et al., 2021; Huijbers et al., 2018; Samire et al., 2023; Sorigué et al., 2017; Zhang et al., 2019) and its detailed mechanism has been unraveled through a wealth of biophysical and biochemical experiments (Sorigué et al., 2021). FAP was first discovered in the green microalga *Chlorella variabilis* NC64A (Sorigué et al. 2017) and was later shown to be conserved with the same activity in a wide variety of algae (Moulin et al. 2021). From a biotechnological standpoint, FAP is interesting because it acts directly on fatty acids and no electron donor or additional cofactor other than the native FAD is required (Santner et al., 2021; Sorigué et al., 2017). Besides, the FAP mechanism does not create a terminal double bound in the HC product compared to other fatty acid decarboxylases such as OleT_JE_ (Jaroensuk et al., 2020) FAP has thus attracted the attention of biotechnologists and chemists. The FAP from *Chlorella variabilis* (*Cv*FAP) and derivative mutants have been used at the laboratory scale not only to produce HCs (Chanquia et al., 2022; Li et al., 2023; Moulin et al., 2019; Ian S. Yunus et al., 2021; Yunus et al., 2018) but also to synthesize high-value chemicals with high yield and enantioselectivity (Emmanuel et al., 2023; Zhang et al., 2020) and perform deracemization of racemic mixtures (Cheng et al., 2020; Li et al., 2021).

The interest in the biotechnological potential of FAP for the production of medium-chain HCs has been recently revived by the demonstration that *Cv*FAP, under specific conditions, can efficiently synthesize C7 and C9 alkanes (Samire et al. 2023). As a purified enzyme, *Cv*FAP displays more activity on octanoic acid (C8:0) than on palmitic acid (C16:0). This increased activity on octanoic acid may be due in part to an autocatalytic effect of the heptane product. Moreover, a faster turnover of the substrate, by reducing FAD triplet formation, may explain the higher stability of FAP observed in the presence of octanoic acid (Wu et al., 2021). It should be noted that the high *in vitro* conversion efficiency of octanoic acid to heptane by FAP is also reflected by feeding experiment; in a bioconversion process where fatty acids are externally supplied to a bacterial culture expressing *Cv*FAP, the conversion rate of octanoic acid to heptane outperforms over 10 times that of palmitic acid to pentadecane (Samire et al., 2023). However, the FAP-based bioproduction of heptane would still be hampered by the absence of a vegetable oil highly enriched in octanoic acid (Jing et al., 2011; Ohlrogge et al., 2018). Thus a sustainable production of heptane by microbial systems entails that the microbes produce octanoic acid. This fatty acid can be synthesized by some plants such as *Cuphea hookeriana*, and bacteria, such as *Annaerococcus tetradius*, which possess medium-chain thioesterases that can preferentially hydrolyze octanoyl-CoA/ACP molecules to octanoic acid (Dehesh et al., 1996; Jing et al., 2011).

In this study, we explore the heptane production potential of an *E. coli* strain engineered to express *Cv*FAP and a thioesterase supplying the octanoic acid precursor. Various FAPs and thioesterases are tested. With a view to practical industrial applications, we use a light-inducible promoter, thus eliminating the need for chemical inducers, and we tested the production capacity for one of the most promising strains in 100-mL photobioreactors. Our study therefore provides a first proof-of-concept of a microbial photoproduction process of heptane and identifies key issues for its future improvement and scale-up.

## 2. Materials and methods

### 2.1. Plasmid construction and strains used

Synthetic genes for the different FAP and octanoyl-ACP thioesterases (C8TES) were codon-optimized for expression in *E. coli*. The sequences of the synthetic genes are detailed in **Table SA**. The FAP genes encode the proteins without their predicted chloroplast transit peptide (Moulin et al., 2021). The FAP gene from *Chlorella variabilis* NC64A (CvFAP) was cloned in pLIC03 and in pLIC07 using the BsaI restriction site. The main difference between these two constructions is the presence of a thioredoxin (TrxA) in-frame with CvFAP in pLIC07. The other FAP genes were also cloned in the pLIC07 plasmid, as described previously (Moulin et al., 2021; Sorigué et al., 2017). For cloning CvFAP in the pDAWN plasmid, the TrxA-CvFAP fragment was amplified by PCR from the pLIC07-CvFAP construct. The product was cloned between HindIII and XhoI restriction sites. For cloning the different C8TES among TrxA-CvFAP in pDAWN plasmid, firstly, the C8TES coding sequence from *Cuphea hookeriana* (ChTES) was cloned in the pDAWN plasmid between NheI and NotI restriction sites, creating the pDAWN-ChTES plasmid. Secondly, the TrxA-CvFAP coding sequence was cloned in the pDAWN-ChTES plasmid between NheI and XhoI. For cloning the remaining C8TES (CpTES-287 or AtTES) coding sequences, the pDAWN-ChTES+CvFAP was digested with NheI and NotI restriction enzymes to remove the ChTES coding sequence and replace it with the CpTES-287 or AtTES coding sequences (See **Table SB**, for constructions and **Table SC** for primers). For heterologous protein expression, we used the following *E. coli* strains: BL21(DE3), Rosetta™(DE3), Rosetta-gami™(DE3), C41(DE3), C43(DE3). The different *E. coli* strains also contain pRIL or pRARE2 plasmids to improve protein expression. (**Table SD)**.

### 2.2. Culture conditions in shake flasks

*E. coli* cells were pre-cultured at 37°C overnight in Luria-Bertani (LB) broth medium with antibiotics (kanamycin 50 µg.mL^-1^ and chloramphenicol 34 µg.mL^-1^). Cultures were initiated at an optical density (OD600) of 0.1 in terrific broth (TB) medium (with antibiotics). All experiments were made in triplicate using 100 ml shake flasks containing 30 ml of culture. Cells were grown in an orbital shaker at 37°C, 200 rpm in the dark. When OD600 reached ∼0.8, the temperature was reduced to 20°C, and protein expression was induced either with different IPTG concentrations (0-2 mM) or different blue-light intensities (0-1.2 µmol.m^-2^.s^-1^) using LEDs. The cultures were grown in these conditions during 16 or 18 h, for experiments with added octanoic acid, after a 16 h induction period, 300 µL of ethanol containing octanoic acid were added to the cultures (0-8 mM final) followed by an incubation period of 2 h in the dark) at 20°C. Samples were then collected for SDS-PAGE, immunoblot analysis, fatty acid analysis byGC-MS/FID (Gas Chromatography coupled to Mass Spectrometry and Flame Ionization Detection) or to determine the heptane production capacity (see below).

### 2.3. Determination of the heptane production capacity

Five ml of cultures from shake flasks were transferred to 10 mL vials which were hermetically sealed before illumination with 300 µmol m^-2^.s^-1^ of blue light for 20 min to 6 h to investigate production within the cells and in the gas phase. To evaluate total HC production, vials were heated at 100°C for 20 min to quench the enzymatic reactions and lyse the cells. Vials were cooled down and kept at 40°C for 5 min. Total HCs were then analyzed via headspace GC-MS/FID. To quantify HCs present within the cells after the illumination period, vials were simply placed in the dark to stop the reaction and volatile HCs were then analyzed via headspace GC-MS/FID. A schematic representation of the experiment is shown in **Figure S12 and S13**. The blue LED spectrum used in this experiment is found in **Figure S14A**.

### 2.4. Analysis of HCs and fatty acids in cells

For the quantifications of HCs and fatty acids present in cells cultivated in shake flasks, samples were normalized on OD600. For each sample, a volume of cell culture equivalent to 1 ml at OD600 10 was pelleted at 3200g for 15 min. When cells were previously incubated with octanoic acid, the pellets were washed and centrifuged 3 times with TB to remove the remaining fatty acids. Internal standards (10 µg of hexadecane and 10 µg of triheptadecanoylglycerol) were added for quantification and cell pellets were transmethylated with 1 ml of methanol containing 5% (v/v) sulfuric acid by heating for 90 min at 85 °C in sealed glass tubes. After cooling down, 1.5 ml of 0.9% (w/v) NaCl and 500 µL of hexane were added. Samples were then shaked for 5 min and centrifuged at 3200 g for 5 min to allow phase separation and recovery of HCs and fatty acid methyl esters (FAMEs) in the organic phase. Finally, the hexane phase is analyzed by GC-MS/FID.

### 2.5. *E. coli* cell viability assays

To assess the toxicity of octanoic acid in *E. coli*, cells from vials added with octanoic acid and illuminated were diluted dilutions (1:10 to 1:10^8^) and 5 µL were dropped on LB 15% (w/v) agar plates. Plates were then kept overnight at 37°C before observation.

### 2.6. SDS-PAGE and immunoblot analysis

One milliliter of *E. coli* shake flask culture was collected by centrifugation at 11,000 g for 5 minutes, followed by resuspension of pellets in 50 mM TRIS buffer pH 8 and sonicated. Cell lysates were mixed with 2X NuPAGE™ LDS Sample Buffer and 50 mM DTT before being boiled for 20 minutes at 70ºC. Proteins were loaded on a constant OD600 basis and separated on SDS PAGE using BisTris 10% precast gels. For protein size determination, the protein ladder PageRuller™ was used and proteins were stained with ProSieve™ EX Safe Stain. For immunoblot analysis, proteins were transferred onto a BioTrace NT nitrocellulose membrane (Sigma-Aldrich). The membrane was blocked with 5% (w/v) dried milk in Tris Buffer Saline containing 0.1% (w/v) Tween 20 overnight at 4°C. For the detection of His-tagged FAPs and homolog proteins, as well as for the detection of His-tagged C8TES proteins, the membrane was incubated at room temperature for 1 h with horseradish peroxidase-conjugated rabbit anti-His antibody coupled to horseradish peroxidase (HRP) (Amersham Biosciences). For detection of CvFAP protein in bacterial strains harboring CvFAP genes together with C8TES genes in pDAWN plasmids, the membrane was incubated with specific primary polyclonal antibodies from rabbit (dilution 1:2000) for 1 h and then for 1 h with a secondary anti-rabbit antibody coupled to HRP (Amersham Biosciences). Immobilon TM Western Chemiluminescent HRP substrate (EMD Millipore) was used for detection. Images were recorded using a G:BOX Chemi XL (Syngene).

### 2.7. Culture of *E. coli* in 100 mL-photobioreactors

To monitor the production of heptane in 100 mL-photobioreactor (turbidostat), bacterial cultures (80 mL) were grown using the Multi-Cultivator MC 1000-OD from Photon Systems Instruments. A blue LED panel combined with neutral density filters (LEE filters) was used to provide light intensity ranging from 0 to 590 µmol.m^-2^.s^-1^. For HC recovery, desorption liners packed with Tenax TA™ (Product number: 020810-005-00, Gerstel, Mülheim an der Ruhr, Germany) were plugged into the air outlet. Antifoam 204 (0.02% final volume) was added to the culture media (TB) to avoid foaming. The initial optical density of the cultures was set at 0.2 from pre-cultures. Cultures were grown at 37°C in darkness with a continuous airflow of 5 L.h^-1^ until OD600 reached 0.8. Various culture conditions were tested, including temperature (from 15°C to 25°C), and airflow (from 1.5 to 5 L.h^-1^). The cultures were then illuminated with LED panels at different blue-light intensities ranging from 0 to 590 µmol photons.m^-2^.s^-1^ for either 18 h or 104 h. For the capture of volatile HCs, a tube containing Tenax TA™ adsorbent resin was inserted in the photobioreactor air outlet for 15 minutes and then transferred to a thermal desorption unit (TDU) coupled with GC-MS/FID analysis for HC quantification. A schematic representation of the photobioreactor system and all the devices equipped is detailed in **Figure 6A** and **Figure S11A**. The blue LED spectrum used in this experiment is found in **Figure S14B**.

### 2.8. GC-MS/FID analyses

For GC-MS/FID analyses, a GC-FID 7890B Agilent coupled to a 5977B series mass detector and equipped with a TDU, a MultiPurpose autosampler (Gerstel) and a cooled injection system (CIS) was used. The MS was run in full scan over 40-350 Th (electron impact ionization at 70 eV), peaks were identified based on their retention time and mass spectrum and quantified based on the FID signal using internal standards or external standard curves.

For analysis of FAMEs and HCs extracted with hexane, we used a HP-5MS Column (30 m by 0.250 mm, 0.25 μm), we injected one µL of the hexane phase into the GC. We used helium (1.4 ml/min) as the carrier gas and liquid N2 was used to decrease the oven temperature. The GC parameters were as follows: oven initial temperature, 5°C for 1 min; ramp, 20°C/min to 160°C and then 10 °C/min to 300°C temperature is maintained for 1 min; splitless.

For Headspace GC-MS/FID analysis, we used a HP-PLOT Q PT column (30 m, 0.32 mm, 20.00 µm). One ml of gas phase (headspace) of sealed vials was collected with a pre-heated syringe (80°C) and injected into the GC. We used Helium as the carrier gas (1.4 mL/min). The GC parameters were as follows: oven initial temperature, 50°C for 1 min; ramp, 20°C/min to 260°C for 20 min. The amount of volatile HCs was calculated *via* a standard calibration curve of heptane. To mimic as much as possible the experimental conditions, the heptane calibration curve was performed in *E. coli* cell cultures, which carried a pDAWN empty vector and were at OD600 = 7.

For the photobioreactors experiments, volatile HCs trapped on tubes were quantified by GC-MS/FID analysis. The TDU was used to release the volatile HCs, which were then condensed by the CIS and focalized on column head. The analysis parameters were as follows: for the TDU, initial temperature, 50°C; ramp, 100°C/min to 280°C for 4 min For the CIS, initial temperature, 50°C; ramp, 10°C/s to 250°C for 4 min For the GC-MS, oven initial temperature, 50°C for 2 min; ramp, 20°C/min to 260°C for 8 min. Column and MS parameters used were the same as in headspace analysis and volatile HCs was calculated *via* an external standard curve obtained by injecting different heptane amounts into the liners packed with Tenax TA™.

## 3. Results and discussion

### 3.1. Effect of thioredoxin fusion on the FAP activity in *E. coli*

In our previous studies on FAP, the recombinant protein was systematically fused to *E. coli* thioredoxin A (TrxA) in the N-terminus because it was initially observed that the TrxA-FAP fusion shows increased solubility during the FAP purification process (Sorigué et al. 2017). However, the benefit of TrxA presence for the activity of FAP in *E. coli* cells was not evaluated. Therefore, to determine if the metabolic burden created by the addition of TrxA was necessary for HC synthesis in *E. coli*, we expressed the most characterized and widely used FAP homolog, *Cv*FAP, in *E. coli* BL21(DE3) cells, fused or not with TrxA, under the control of the IPTG-inducible P_T7_ promoter (**Table A, SA-B & Figure 1A-B**). The results clearly show that *Cv*FAP fused with TrxA resulted in a 12-fold and a 8-fold increase in total HC content at 24 h, and 48 h post-induction respectively (**Figure 1C**) while cell concentrations were similar with and without TrxA **(Figure S1A)**. Although TrxA fusion slightly changed the HC profile, the species produced remained mainly C15-C17 alka(e)nes (**Figure S1B**). Protein and immunoblot analysis of total and soluble fractions indicated that the expression level of CvFAP fused with TrxA was much higher after 24 and 48 h of induction with IPTG compared to CvFAP alone (**Figure 1D, Figure S2)**. This observation suggested that TrxA also helped solubility and/or stability of FAP *in vivo*.

**Figure 1:**
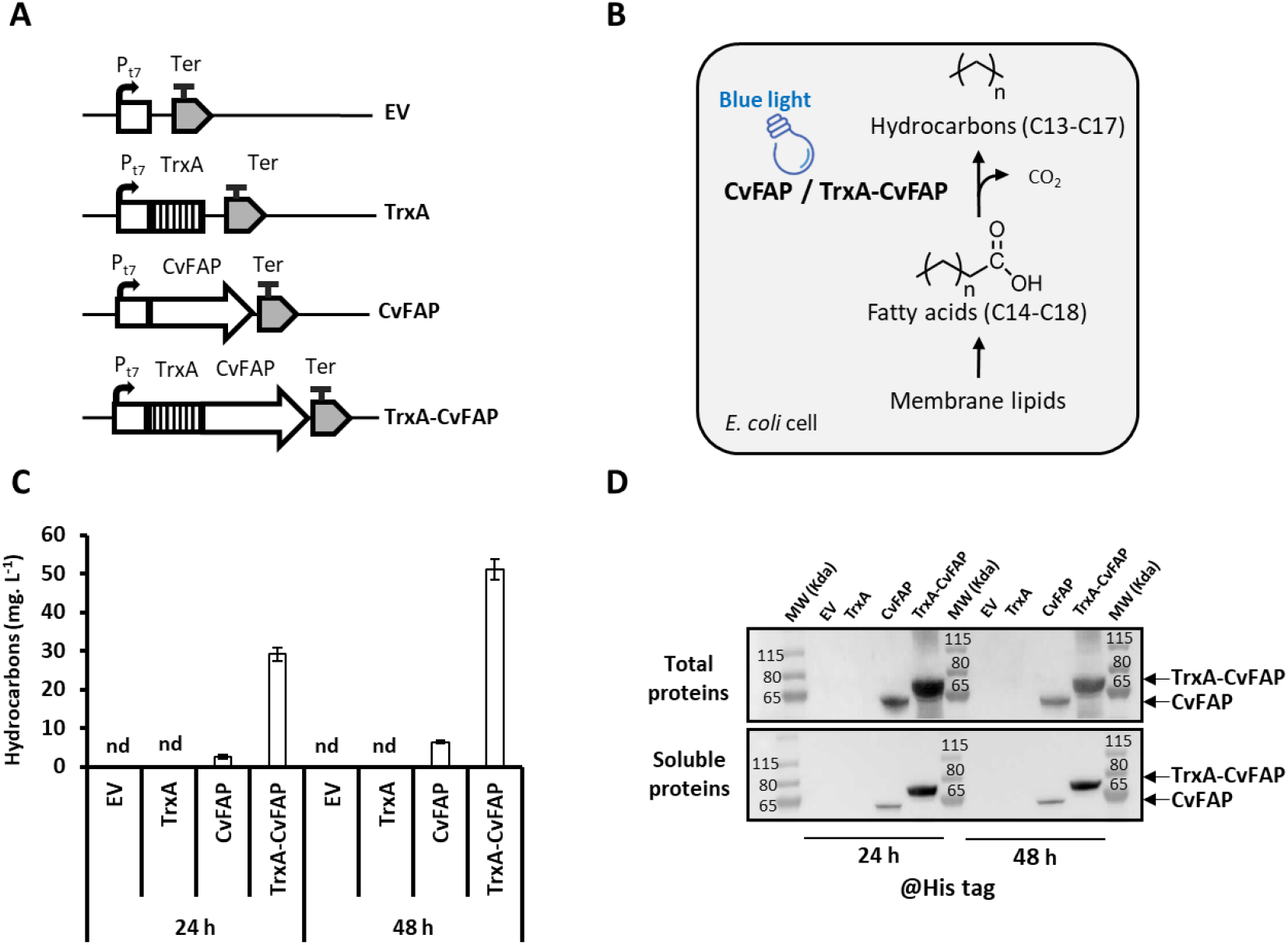
Impact of TrxA Fusion to the N-terminus of CvFAP on Hydrocarbon Production in *E*.*coli* BL21(DE3). **A)** Scheme of the genetic constructs employed. Pt7 = T7 promoter, TrxA = *E. coli* thioredoxin A, Ter = terminator, EV = empty vector. **B)** Schematic representation of metabolic pathways involved in hydrocarbon production. **C)** Hydrocarbons detected *in E. coli* cells expressing CvFAP with and without TrxA fusion. Cells were induced with 0.5 mM IPTG and cultured in shake flasks under 100 µmoles photons.m^-2^.s^-1^ of blue light for 24 and 48 hours. **D)** Immunoblot analysis of CvFAP in the total and soluble protein fractions of *E. coli* cells collected at 24 and 48 hours post-induction. The full immunoblot and corresponding protein gel are shown in Figure S2. Protein loading was normalized based on cell density (OD600). Error bars represent the standard error of three biological replicates. ‘nd’ = not detected.

We also observed that TrxA-FAP, but not TrXA alone, increased the total fatty acid content with a more significant effect at 48h **(Figure S1C)**. The presence of CvFAP also changed drastically the fatty acid profile by increasing the amount of C18:1 fatty acid and decreasing pentadecanoic and cyclo pentadecanoic acid significantly compared to control strains **(Figure S1D)**. This indicated a metabolic adaptation of the host to compensate for fatty acid depletion and/or conversion into hydrocarbons caused primarily by CvFAP.

To what extent TrxA modifies FAP turnover rate or stability in addition to increasing solubility remains to be established. However, the overall positive effect on the total heptane production clearly shows that TrxA fusion should be used in any biocatalytic or bioconversion process involving *Cv*FAP. The substantial enhancement in CvFAP expression and hydrocarbon production due to TrxA fusion led us to systematically employ TrxA-FAP fusions in all subsequent experiments.

**Table A:**
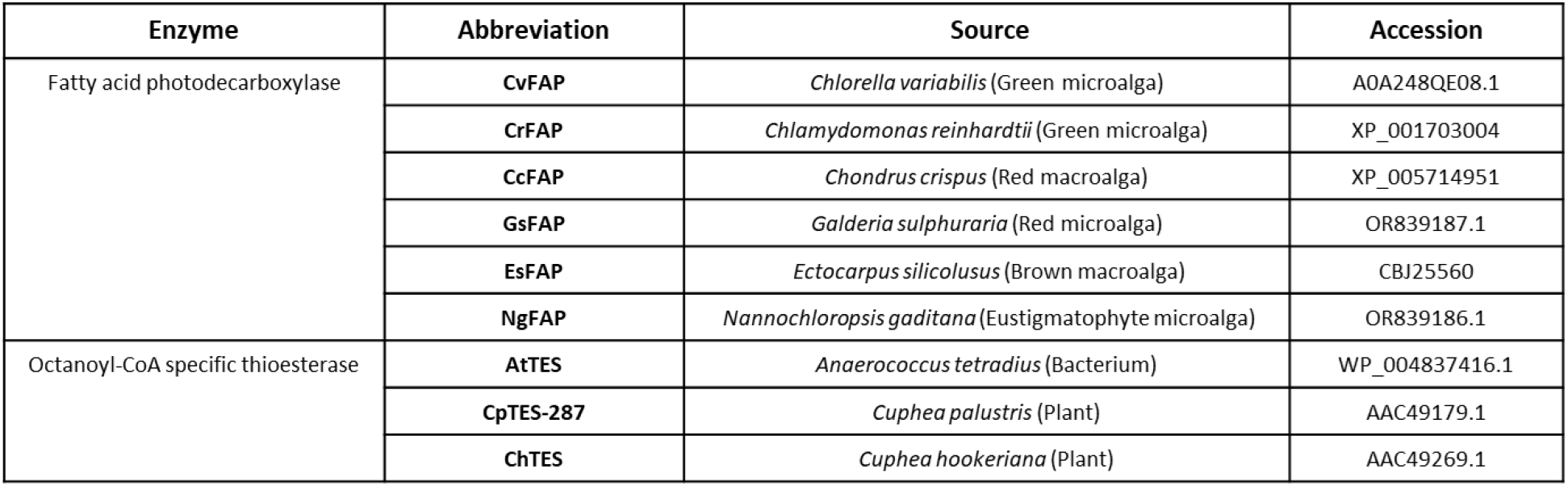
List of Fatty Acid Photodecarboxylases (FAPs) and Octanoyl-CoA Specific Thioesterases (C8_TESs) Used in This Study. The amino acid sequence of CpTES-287 has been modified from its original sequence. It features a truncation of 112 residues from the N-terminus and includes two point mutations: N122S and I159M. This variant was originally designated as CpFatB1.2-M4–287 (Hernández Lozada et al., 2018). The *Ch*TES variant was originally designated as *Ch*FatB2 (Dehesh et al., 1996).

### 3.2. Test of FAP homologs for heptane production

To date, only *Cv*FAP has been documented to produce heptane (Samire et al., 2023; Yunus et al., 2021). Exploring the conservation of FAP activity in various microalgae, a few other homologs with demonstrated photodecarboxylase activity were identified (Ma et al., 2023; Moulin et al., 2021; Zeng et al., 2022). These studies suggested that some homologs may have a different substrate specificity profile than *Cv*FAP but activity on octanoic acid was not tested. We therefore assessed the capacity of five of these FAPs for heptane production. To simplify the screening process, octanoic acid was directly added to the culture media. We first tested the effect of octanoic acid on cells by adding different concentrations of octanoic acid to *E. coli* cells harboring an empty vector. We observed that octanoic acid concentration under 4 mM did not affect cell viability (**Figure S3A**) and therefore decided to use 2 mM of octanoic acid. A first series of experiments indicated that in the empty vector strain grown for 16 h in the dark, octanoic acid could be detected inside cells as early as 2 h after addition to the medium (**Figure 2A**). FAP homologs were expressed under the control of the IPTG-inducible P_T7_ promoter (**Figure 2B-C**), SDS-PAGE and immunoblot analyses indicated that all FAPs were well-expressed. In the soluble fraction only *Cv*FAP, *Cr*FAP, *Cc*FAP and *Ng*FAP were detected, CvFAP being the most abundant (**Figure 2D, Figure S3B-E**). Growth of *E. coli* strains was slightly affected by expression of FAPs except for GsFAP (**Figure S3F**). To measure the heptane production capacity of each strain expressing a FAP, the shake flask cultures pre-incubated with octanoic acid were exposed for 0.33 h to 300 µmol.m^-2^.s^-1^ of blue light in sealed vials and heptane was measured in the gas phase. All FAPs homologs tested were found to be active on octanoic acid but the heptane levels varied a lot and the highest production of heptane was obtained with *Cv*FAP (**Figure 2E**). The fact that *Cv*FAP (fused to TrxA) outperforms other FAP homologs (also fused to TrxA) for heptane production *in vivo* could be due in part to the lower amounts of FAP soluble form **(Figure 2D)** but differences between FAPs in turnover on octanoic acid and in stability cannot be ruled out. In practice, the strain harboring *Cv*FAP had clearly the best capacity to convert octanoic acid to heptane and it was therefore selected for the rest of the work.

**Figure 2:**
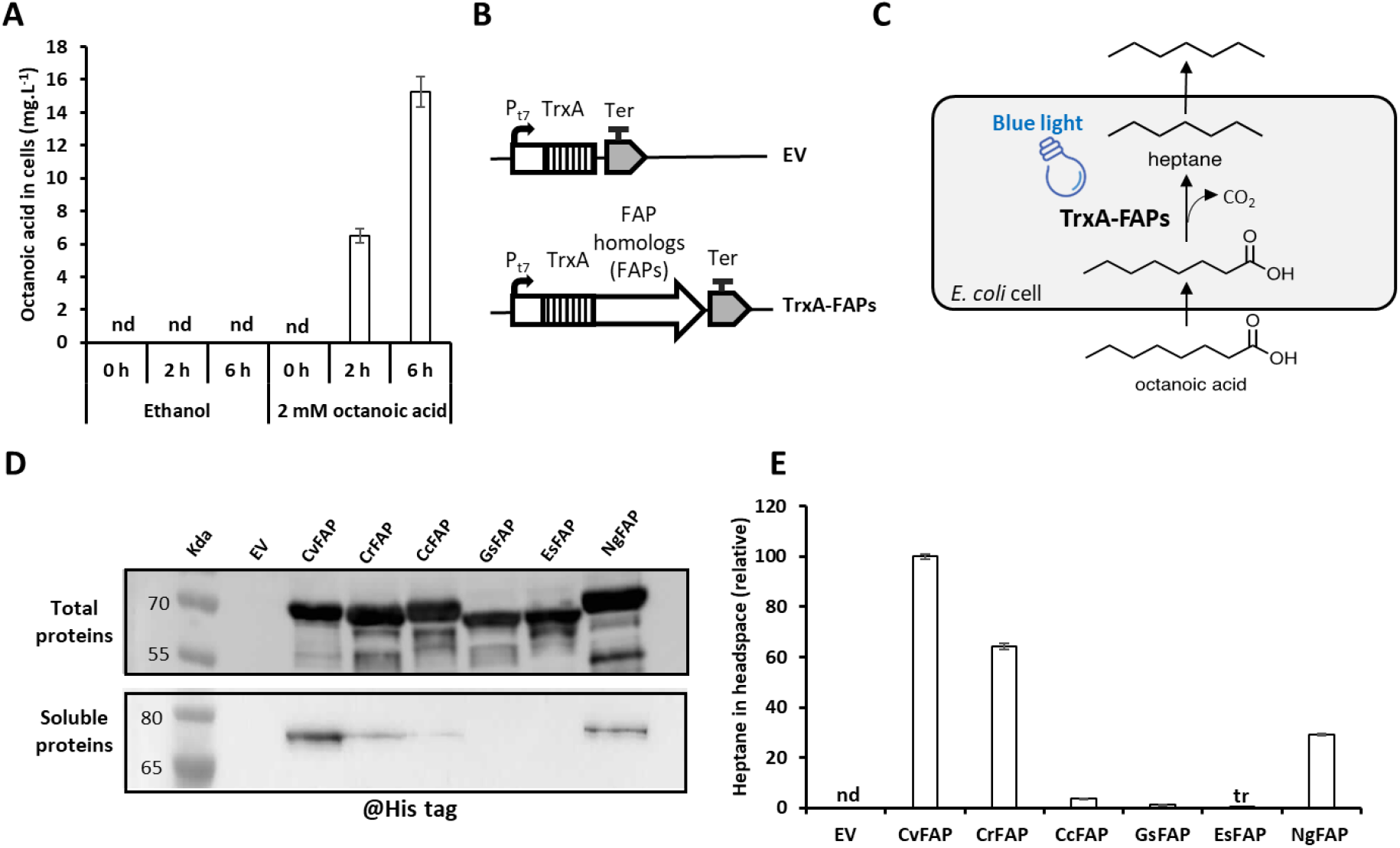
Heptane Production in the Gas Phase of *E. coli* Cultures Expressing Various FAPs under Blue Light Illumination with Octanoic Acid Supplementation. **A)** Time course of octanoic acid uptake by E. coli cells harboring an empty vector (EV), treated with 2 mM octanoic acid (in ethanol) after 16 h of growth. Cells were washed twice before quantification. *E. coli* strains exposed only to ethanol served as controls. **B)** Schemes of the genetic constructs used in this study. **C)** Schematic representations of the metabolic pathways. **D)** Immunoblot detection of various FAP homologs fused to TrxA in the total and soluble protein fractions from E. coli strains after a 16-hour culture period in the dark. Protein loading was normalized based on cell density (OD600). **E)** Relative quantification of heptane production in the gas phase of *E. coli* cultures expressing different FAP homologs fused to TrxA, compared to control cultures with an EV. After growing in the dark for 16 hours, cultures were treated with 2 mM octanoic acid for 2 hours and then incubated under 300 μmol photons m^-2^ s^-1^ of blue light in sealed vials for 0.33 hours at 20°C. Error bars represent the standard error based on three biological replicates. The full immunoblot and protein gel are presented in Figure S3. ‘nd’= not detected, ‘tr’= trace.

### 3.3. Comparison of IPTG and blue light-inducible promoter for heptane production

To streamline a bioprocess and eliminate the need for chemical inducers, we chose to investigate the use of a light-inducible promoter for controlling *Cv*FAP expression. We selected the blue light-inducible promoter P_DAWN_ (Ohlendorf et al., 2012) because blue light also triggers FAP activity. In other words, with P_DAWN_, the blue light illumination used during the HC production phase may thus also serve to facilitate the constant replacement of FAP, which is prone to photoinactivation (Lakavath et al., 2020).

We first compared the performances of P_DAWN_ with IPTG inducible promoter P_T7_ to control the expression of *Cv*FAP (**Figure 3A)**. *E. coli* strains were cultured for 16 h, and gene expression was induced under different light intensities ranging from 0 to 1.2 μmol photons.m^-2^.s^-1^ or IPTG concentrations (0-2 mM). The cell growth decreased with increasing concentrations of IPTG, and it remained constant with increasing light intensities (**Figure S4**). Interestingly comparing the heptane production in the *E. coli* strain expressing *Cv*FAP under P_DAWN_ or P_T7,_ we found that the use of IPTG resulted only in a 20% higher heptane production compared to light, (**Figure 3B**), demonstrating that the P_T7_ promoter can be advantageously replaced by the P_DAWN_ promoter for *Cv*FAP expression. Using P_DAWN_, we showed that a 0.2 µmol.m^-2^.s^-1^ allows the highest heptane production and we selected this induction condition. Thanks to its low light intensity requirement, only a very low amount of energy is needed to provide the proper blue light intensity to induce P_DAWN_, which may help avoid photoinactivation of FAP during the induction phase. Sunlight may also be used to activate the promoter. We also tested the effect of octanoic acid on *E. coli* cells expressing *Cv*FAP under the control of P_DAWN_ promoter. We observed that even 8 mM of octanoic acid did not affect cell viability (**Figure S4B**). It seems that the conversion of octanoic acid to heptane by *Cv*FAP nullifies the toxic effect of added and produced octanoic acid.

**Figure 3:**
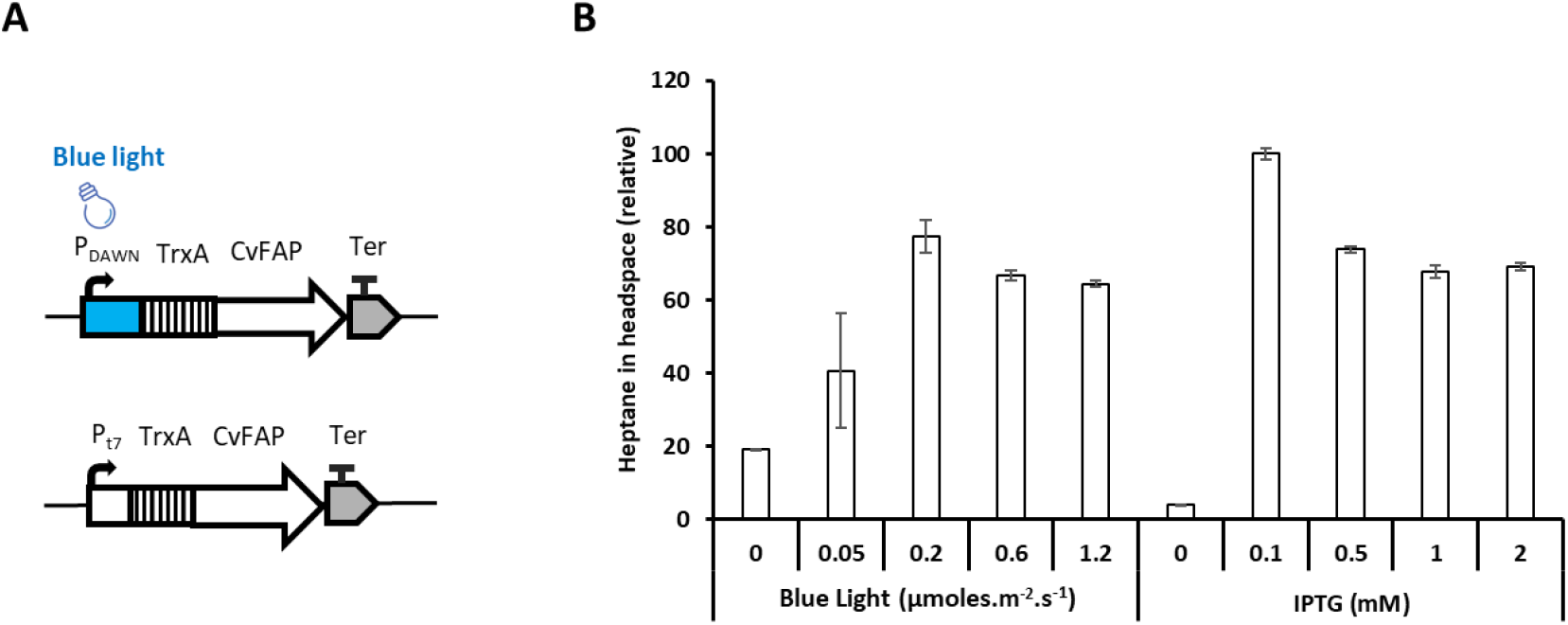
Comparison of HC production of BL21(DE3) E.coli strains expressing CvFAP under the control of Blue Light-Inducible and IPTG-inducible promoters. **A)** Schemes of the genetic constructs used, featuring both blue light-inducible (P_DAWN_) and IPTG-inducible (Pt7) promoters. **B)** Relative quantification of heptane produced in the gas phase of *E. coli* cultures expressing *Cv*FAP under either P_DAWN_ or Pt7. The strains expressed *Cv*FAP under the control of either P_DAWN_ or Pt7 and were exposed to varying light intensities (0-1.2 μmoles.m^-2^.s^-1^) or different IPTG concentrations (0-2 mM), respectively during 16 h. Then, cultures were incubated with 2 mM octanoic acid for 2 hours in darkness, then exposed to 300 μmol photons m^-2^s^-1^ of blue light in sealed vials for a 0.33 hours period at 20°C. Error bars represent the standard error based on five biological replicates.

### 3.4. Selection of an octanoyl-ACP-specific thioesterase for endogenous substrate supply

To build an *E. coli* FAP-expressing strain that produces its own octanoic acid we evaluated several octanoic acid-producing thioesterases (C8TESs) from plant or bacterium reported in the literature and we also used an improved version of C8TES from *Cuphea palustris* (Hernández Lozada et al., 2018) (**Table A** and **Table SA**). C8TESs interrupt the fatty acid elongation pathway by specifically hydrolyzing octanoyl-ACP intermediates, thereby releasing octanoic acid within the cell (**Figure 4A and S5**). The thioesterase sequences were codon-optimized for *E. coli* expression. The co-expression of C8TES and CvFAP genes was performed in operon under the control of the P_DAWN_ promoter (**Figure 4B**). Protein expression was induced by 0.2 µmol photons.m^-2^.s^-1^ of blue light. We found that the growth of *E. coli* strains was only slightly affected by co-expression of *Cv*FAP and different C8TES compared with the strain expressing *Cv*FAP only (**Figure S6**). As expected, coexpression of each of the thioesterases with *Cv*FAP resulted in the formation of octanoic acid (**Figure 4C**, 18h), with the thioesterase from the plant *Cuphea hookeriana (Ch*TES) being the most producing enzyme, even with a very low expression level (**Figure S7A-B**).

**Figure 4:**
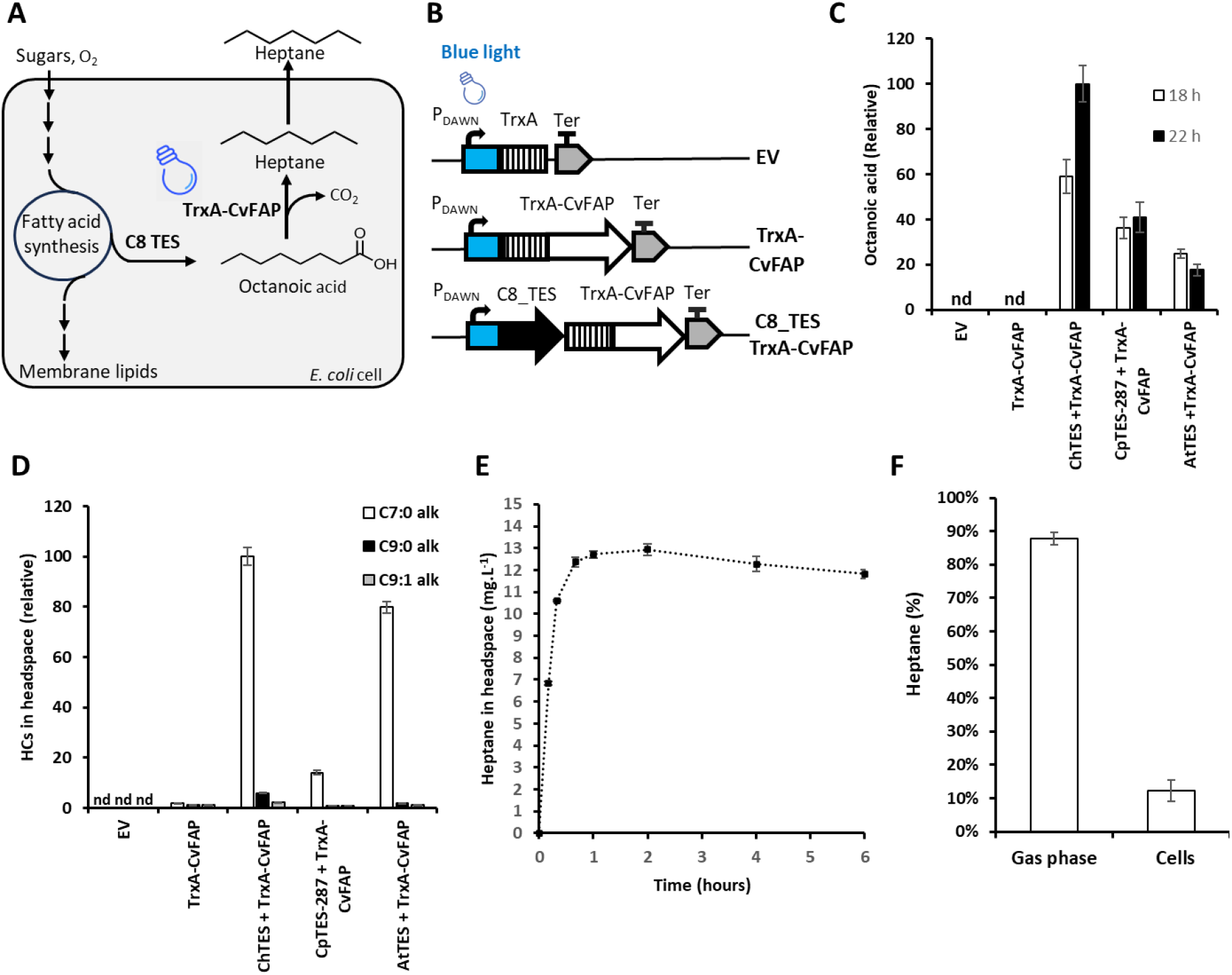
Co-expression of Various Octanoyl-ACP Substrate-Specific Thioesterases (C8TES) and CvFAP Genes Under Blue-Light Inducible Promoter in *E. coli* BL21(DE3). **A)** Schematic representation of the metabolic pathways involved in heptane formation. **B)** Scheme of the genetic constructs. **C)** Octanoic acid content of *E. coli* cells co-expressing *Cv*FAP and different His-tagged C8TES before (18 h) and after (24 h) the illumination of the cultures. **D)** Volatile hydrocarbon levels measured in the headspace produced by *E. coli* cells co-expressing ChTES and CvFAP genes, following 4 hours of incubation under 300 μmol photons.m^-2^.s^-1^. **E)** Kinetics of heptane production in *E. coli* strains co-expressing His-tagged *Ch*TES and *Cv*FAP. Heptane was measured in the headspace of vials at various time points of the illumination at 300 μmol photons.m^-2^.s^-1^. **F)** Proportion of heptane present in the headspace and inside cells produced by *E. coli* cells co-expressing the His-tagged *Ch*TES and *Cv*FAP genes after 4 h of incubation at 300 μmol photons.m^-2^.s^-**1**^. Error bars represent the standard error based on three biological replicates. ‘nd’ = non detected.

Cultures were then exposed to 300 μmol photons.m^-2^.s^-1^ in sealed vials for 4 hours to trigger HC production. Expression of C8TES increased drastically the total HC content in the gas phase of the vial and the highest amount was obtained with *Ch*TES **(Figure 4D)**. A kinetics of heptane production performed on the *Ch*TES+*Cv*FAP strain indicated that after 2 hours of illumination heptane synthesis stopped, reaching a titer of around 12.5 mg per liter of culture **(Figure 4E)**. Composition of the HCs collected in the gas phase were similar for the three C8TES, with over 90% of heptane, the rest being nonane and nonene (**Figure S7C**). We also found that heptane was present inside cells together with longer HCs, mainly pentadecane and heptadecene (**Figure S9A-B**). When assessing the proportion of heptane in the gas phase and inside cells in the *Ch*TES+*Cv*FAP strain, it was found that almost 90% of heptane was in the gas phase (**Figure 4F**). In an attempt to further increase heptane production, we tested the *Ch*TES+*Cv*FAP construct in other *E. coli* expression strains; however, the six other strains tested proved to be equal or inferior to the BL21(DE3) pRIL strain for heptane production (**Figure S9C**).

From these first experiments, it was clear that strains with different *C8TES* did not have the same ability to produce heptane and in view to future optimization, it was important to understand whether there could be reasons other than the level of thioesterase activity. Cell concentration after induction was only slightly higher in the three C8TES strains **(Figure S6)**. Also, there was no significant effect of the C8-specific thioesterases on the total amount of fatty acids **(Figure S8A)**, which is contrasting the enhancement in fatty acid synthesis observed with the expression of a plant C12-specific TES in *E. coli* (Moulin et al. 2019). Regarding the precursor of heptane, it was observed that before illumination octanoic acid represented 5 to 7.4 % of total fatty acids in the three strains and 3.5 to 11% after illumination (**Figure S8B-C**). Interestingly, the *Ch*TES was the most effective for octanoic acid production both before and after the illumination phase and was the only C8TES for which octanoic acid content was higher after illumination than before **(Figure 4C)**. Another factor that may explain the differences in heptane production is clearly the unequal level of expression of *Cv*FAP in the three strains (**Figure S7B**). Results thus indicated that the heptane level was likely to be the result of a complex interplay between FAP and C8TES expression.

To test whether the octanoic acid level could be limiting heptane formation in the best strain obtained so far (*Ch*TES *+ Cv*FAP), post-induction cells were incubated in the dark for 2 h with different concentrations of octanoic acid (0-6 mM) before illumination at 300 μmol photons.m^-2^.s^-1^. While the addition of exogenous octanoic acid did not have any significant effect on cell viability (**Figure S10**), we observed a clear increase in the quantity of heptane (**Figure 5**), which indicated that the octanoic acid was indeed limiting in the *Ch*TES + *Cv*FAP strain. This idea could be contradictory with the accumulation of octanoic acid over time in this strain (**Figure 4C**). This may be explained by considering the dynamics of fatty acid production and the photoinhibition of FAP (Moulin et al. 2019; Lakavath et al. 2020). Adding a large quantity of exogenous octanoic acid may protect FAP from photoinhibition by avoiding FAD triplet formation, leading to higher alkane production, as proposed for in vitro FAP activity (Samire et al. 2023). Although FAP is supposed to be continually renewed by promoter induction, the synthesis of FAP may not compensate completely for its photodegradation. Over time, as FAP degrades due to photoinhibition, octanoic acid production persists through *Ch*TES and may explain octanoic acid accumulation. Identification of FAP mutants less prone to photoinhibition is an important goal for further increasing HC production.

**Figure 5:**
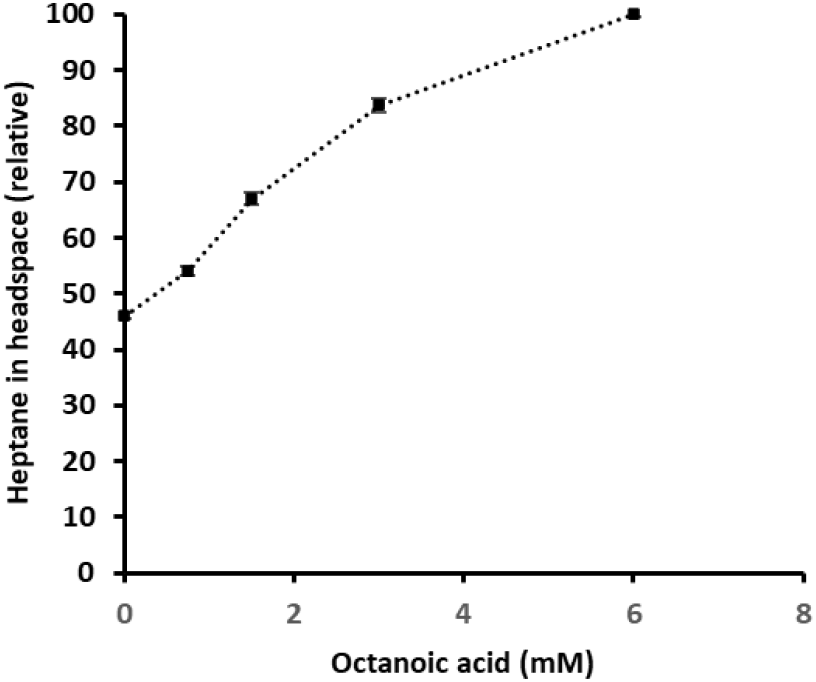
Detection of heptane in *E. coli* strains co-expressing ChTES and CvFAP genes, fed with varying concentrations of octanoic acid and incubated under 300 μmol photons.m^-2^.s^-1^ of blue light in sealed vials for 4 hours at 20°C. Error bars represent the standard error based on three biological replicates.

**Figure 6:**
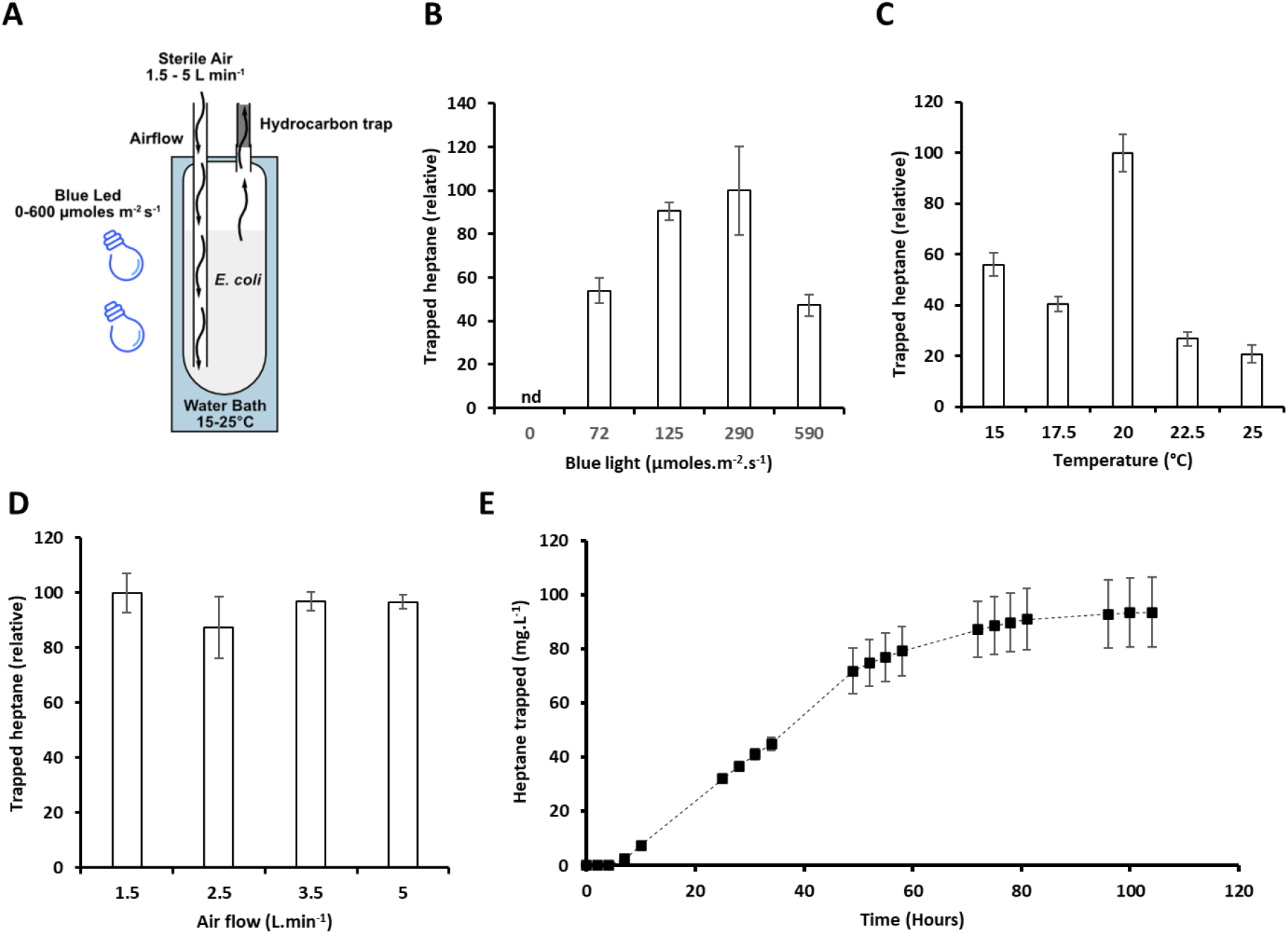
Optimization of Continuous Photoproduction of Heptane in *E*.*coli* BL21(DE3) Co-expressing *Ch*TES and *Cv*FAP in a 100 ml Photobioreactor. **A)** Schematic of the photobioreactor system utilized for heptane production. This system includes adjustable settings for light intensity, temperature, and airflow, and is equipped with desorption liners packed with Tenax TA™, which retains volatile compounds including medium-chain hydrocarbons like heptane. **B)** Relative heptane production from *E. coli* strains cultured for 18 hours at 20°C, with an airflow of 5 L/min, across various blue light intensities (0-590 µmol photons.m^-2^.s^-1^). **C)** Relative heptane production from strains cultured under 290 µmol photons.m^-2^.s^-1^ of blue light exposure for 18 hours at different temperatures. **D)** Relative heptane production from strains cultured under similar conditions with varying airflow rates in the photobioreactor. **E)** Time course of cumulative heptane production in *E. coli* grown under continuous blue light illumination (290 µmol photons.m^-2^.s^-1^) for 104 hours at 20°C and an airflow rate of 1.5 L/min. Error bars represent the standard error from multiple biological replicates (parts B and E n=6, part C n=7, part D n=4). nd = not detected. Relative values were obtained by normalizing to the maximum value measured.

### 3.5. Optimizing photoproduction of heptane in 100 mL-photobioreactors

Studies on the microbial production of HCs were so far mostly explored on a few milliliter scale (Jaroensuk et al. 2020; Geng et al. 2023). In order to evaluate the capacity of the *Ch*TES + *Cv*FAP strain to produce heptane on a larger scale, we used a 100 mL-photobioreactor **(Figure 6A and S11A)**. We investigated the influence of different parameters on hydrocarbon production in the gas phase, including light intensity, temperature and airflow. Growth rates were relatively consistent across the tested conditions, except for the airflow rates, for which higher OD600 values were associated with increased airflow due to media evaporation (**Figure S11B-E)**. The highest heptane production was achieved at 20°C with 290 µmol photons.m^-2^.s^-1^ of blue light (**Figure 6B-D**). Since we observed that higher airflow had no effect on heptane production (**Figure S11E**), we selected the lowest airflow rate of 1.5 L.min^-1^ to limit media evaporation. We grew the strain in these optimized conditions for 104 h. The growth followed a classical bacterial growth curve (**Figure S11F**). Cumulative heptane production reached 94 mg.L^-1^ after 104 h of culture (**Figure 6E**), i.e. an overall productivity of 22 mg.L^-1^.day^-1^. The heptane productivity per hour increased linearly during the exponential phase (up to 10 hours), reaching a maximum of 1.8 mg.L^-1^.h^-1^, remaining stable during the deceleration phases (up to 49 h), then the productivity decreased during the stationary phase (after 49 h), and almost stopped after 100 h, reaching a minimum value of 0.07 mg.L^-1^.h^-1^ (**Figure S11F-G**). These results indicated that heptane production mainly occurred during the exponential and deceleration phase, while significantly decreasing during the stationary phase.

In vial experiments, heptane represented >90% of total HCs in the gas phase (**Figure 4F**) and the titer reached was ∼12.5 mg.L^-1^ of *E. coli* culture in about one hour (**Figure 4E**). In photobioreactors, 22 mg.L^-1^.day^-1^ is equal to 0.9 mg.L^-1^.h^-1^. This indicated a quite strong reduction (>10-fold) in productivity in a 100-fold upscaling. By comparison, when medium-chain saturated HCs, including heptane, were produced by *Synechocystis* sp. PCC 6803 co-expressing the native C8TES from *Cuphea palustris* together with *Cv*FAP in a 10-day culture period (Yunus et al., 2022), the heptane produced at the scale of 25 mL represented ∼5 mg.L^-1^ of culture in 10 days, meaning 0.02 mg.L^-1^.h^-1^. Moreover, heptane was mixed with other medium and long-chain hydrocarbons, including nonane (∼0.5 mg.L^-1^ of culture), undecane (∼0.5 mg.L^-1^ of culture), and tridecane (∼3 mg.L^-1^ of culture) (Yunus et al., 2022). In our experimental setup, we obtained a cumulative heptane production 94 mg per L of culture in 100-mL photobioreactor after 104 h of culture (Figure 6E). Where the productivity achieved a maximum of 1.8 mg.L^-1^. h^-1^ of culture and an overal of 0.9 mg.L^-1^.h^-1^ (45 times increased compared to Yunus and coworkers). Considering this production of medium-chain saturated linear HCs (C5 to C11) in 100 mL-photobioreactor, the yield observed is only surpassed by Choi and Lee’s engineered *E. coli* strain using the plant HC-forming enzyme ECERIFERUM1, which was able to produce 328 mg.L^-1^ of culture of nonane (C9 HC) among other longer chain HCs including dodecane, tridecane, 2-methyl-dodecane and tetradecane (Choi and Lee, 2013). However, the cultivation time necessary to reach this yield were not specified and thus and the strain productivity cannot be calculated.

## 4. Conclusion

In this work, we developed a microbial platform for heptane photoproduction. We fused CvFAP to TrxA and increased its biological activity. We tested different FAP homologs and identified *Cv*FAP as the best enzyme for *in vivo* heptane synthesis. To produce the CvFAP substrate *in situ*, we co-expressed the CvFAP with different thioesterases. We reached a yield of 94 mg.L-1 by co-expressing CvFAP together with the *Cuphea hookeriana* thioesterase in 100-mL photobioreactor. Our work provides a fundamental framework for microbial synthesis of medium-chain hydrocarbons and will hopefully contribute to more sustainable fuel production that is less dependent on fossil resources.

## Supporting information

mains and SD Figures

## CRediT authorship contribution statement

All authors discussed the results and contributed to the final manuscript. A.B.P, B.L., P.A.T., F.V., S.C., P.P.S. & S.M. carried out the experiments. A.B.P., Y.L.B, B.L., F.B. & D.S. contributed to the design and implementation of the research, to the analysis of the results, and the writing of the manuscript.

## Acknowledgements

We thank Dr. Laurence Blanchard for helpful discussions and Dr. Gilles Peltier. The funding provided by the European Union Regional Developing (ERDF), the Région Provence Alpes Côte d’Azur, the French Ministry of Research, and the CEA to the HelioBiotec platform is also acknowledged. A.B.P. was supported by a PhD fellowship “Economie Circulaire du Carbone” from CEA. D.S.’s work was supported by CEA and by a Marie Sklodowska-Curie Actions global fellowship (Project 101063280 - EXPERIMENTAL).

