## Supplementary material for "Microbial photoproduction of heptane": mains and SD Figures

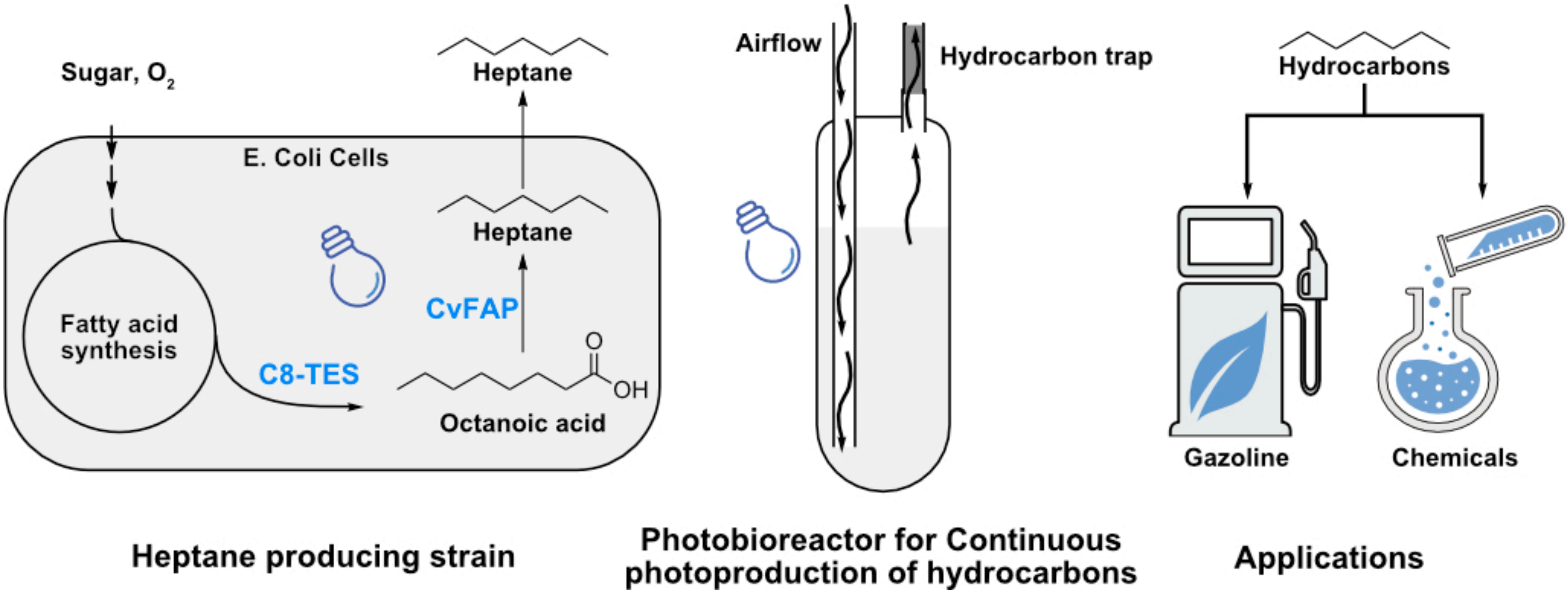
Sugar, $O_2$
E. Coli Cells
Heptane
Heptane
CvFAP
Fatty acid synthesis
C8-TES
O
OH
Octanoic acid
Airflow
Hydrocarbon trap
Hydrocarbons
Gazoline
Chemicals
Heptane producing strain
Photobioreactor for Continuous photoproduction of hydrocarbons
Applications

| Enzyme | Abbreviation | Source | Accession |
|---|---|---|---|
| Fatty acid photodecarboxylase | **CvFAP** | *Chlorella variabilis* (Green microalga) | A0A248QE08.1 |
| | **CrFAP** | *Chlamydomonas reinhardtii* (Green microalga) | XP_001703004 |
| | **CcFAP** | *Chondrus crispus* (Red macroalga) | XP_005714951 |
| | **GsFAP** | *Galderia sulphuraria* (Red microalga) | OR839187.1 |
| | **EsFAP** | *Ectocarpus silicolusus* (Brown macroalga) | CBJ25560 |
| | **NgFAP** | *Nannochloropsis gaditana* (Eustigmatophyte microalga) | OR839186.1 |
| Octanoyl-CoA specific thioesterase | **AtTES** | *Anaerococcus tetradius* (Bacterium) | WP_004837416.1 |
| | **CpTES-287** | *Cuphea palustris* (Plant) | AAC49179.1 |
| | **ChTES** | *Cuphea hookeriana* (Plant) | AAC49269.1 |

**Table A: List of Fatty Acid Photodecarboxylases (FAPs) and Octanoyl-CoA Specific Thioesterases (C8_TESs) Used in This Study.** The amino acid sequence of CpTES-287 has been modified from its original sequence. It features a truncation of 112 residues from the N-terminus and includes two point mutations: N122S and I159M. This variant was originally designated as CpFatB1.2-M4–287 (Hernández Lozada et al., 2018).

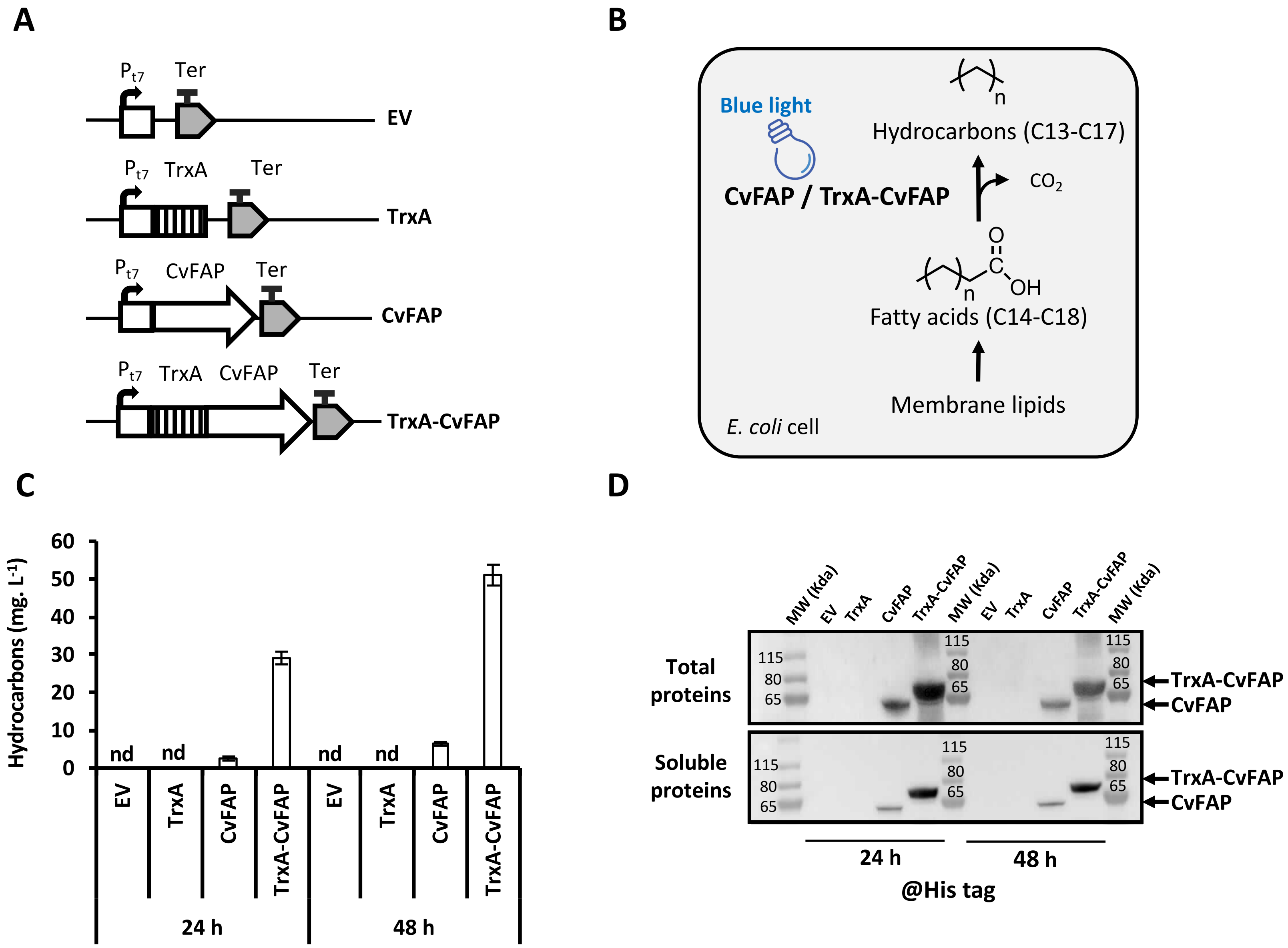


<u>**Figure 1:**</u> **Impact of TrxA Fusion to the N-terminus of CvFAP on Hydrocarbon Production in *E.coli* BL21(DE3). A)** Scheme of the genetic constructs employed. Pt7 = T7 promoter, TrxA = *E. coli* thioredoxin A, Ter = terminator, EV = empty vector. **B)** Schematic representation of metabolic pathways involved in hydrocarbon production. **C)** Hydrocarbons detected *in E. coli* cells expressing CvFAP with and without TrxA fusion. Cells were induced with 0.5 mM IPTG and cultured in shake flasks under 100 µmoles photons.m$^{-2}$.s$^{-1}$ of blue light for 24 and 48 hours. **D)** Immunoblot analysis of CvFAP in the total and soluble protein fractions of *E. coli* cells collected at 24 and 48 hours post-induction. The full immunoblot and corresponding protein gel are shown in Figure S2. Protein loading was normalized based on cell density (OD600). Error bars represent the standard error of three biological replicates. 'nd' = not detected.

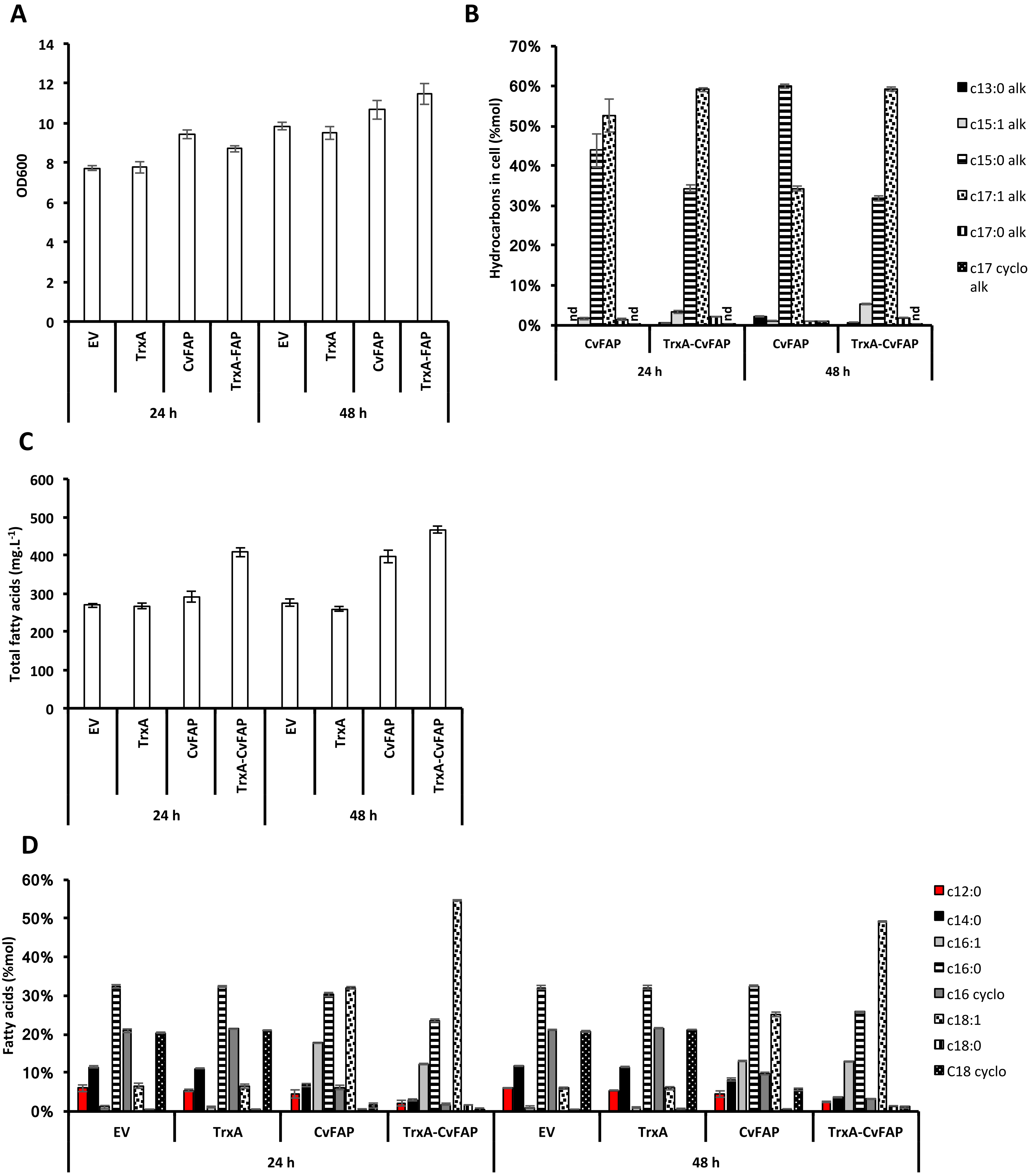


**Figure S1: Influence of TrxA Fusion to the N-terminus of CvFAP on Hydrocarbon and Fatty Acid Content in *E. coli* BL21(DE3) Expressing CvFAP. A)** Optical density (OD600) of *E. coli* cultures expressing CvFAP, with or without TrxA fusion. **B)** Distribution of different hydrocarbons in *E. coli* cells expressing CvFAP with or without TrxA fusion. **C)** Total fatty acid content in E. coli cells expressing CvFAP, with or without TrxA fusion, after induction with 0.5 mM IPTG and exposure to 100 µmol photons·$m^{-2}s^{-1}$ of blue light for 24 and 48 hours. **D)** Distribution of different fatty acids in E. coli cells expressing CvFAP with or without TrxA fusion. Error bars represent the standard deviation based on three biological replicates. . 'nd' = not detected.

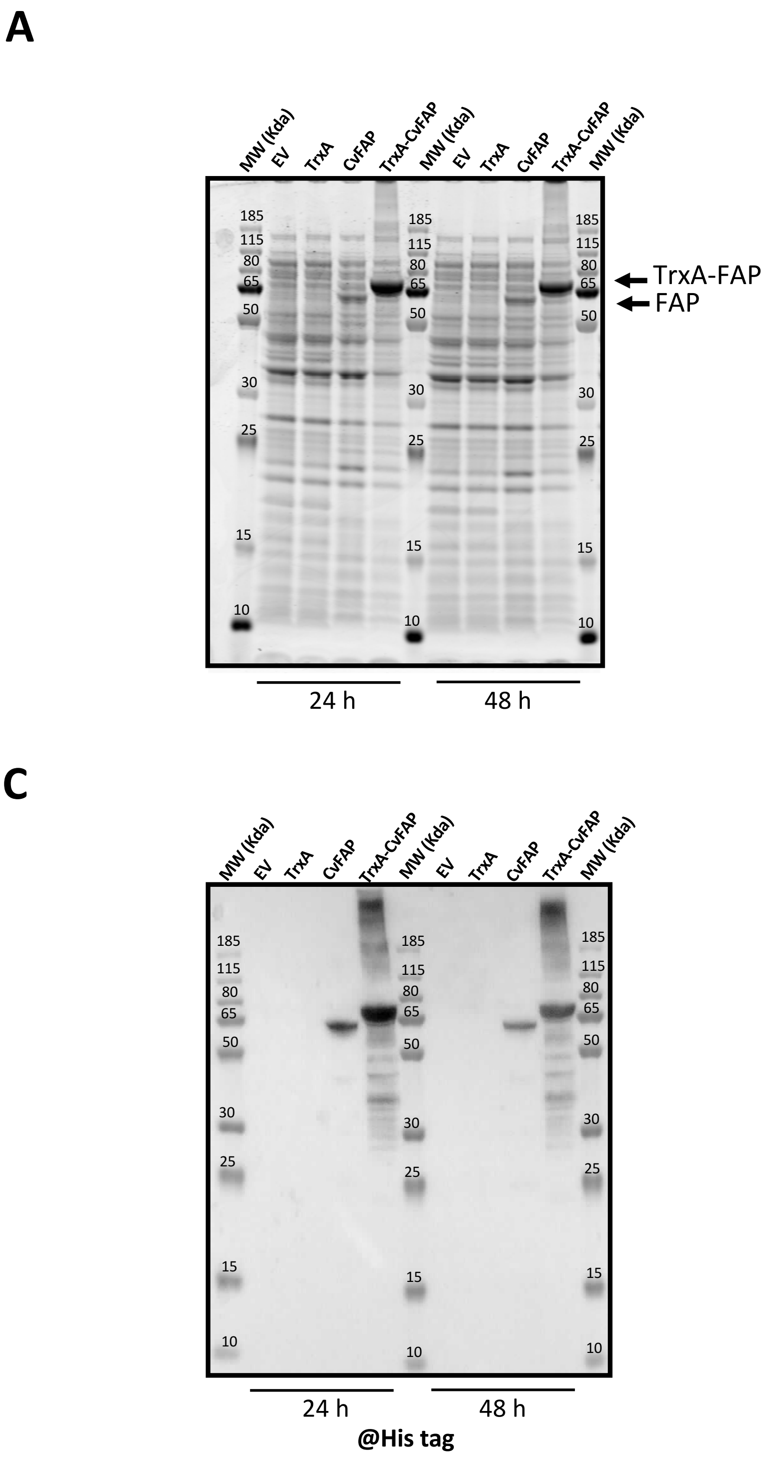


**<u>Figure S2:</u> Analysis of CvFAP Co-expression with and without N-terminal Fusion to *E. coli* Thioredoxin A (TrxA) in *E. coli* BL21(DE3).** SDS-PAGE analysis of total proteins (**A**) and soluble proteins (**B**) from E. coli strains co-expressing CvFAP, with or without TrxA fusion. Immunoblot detection of CvFAP in total protein extracts (**C**) and soluble protein (**D**) of E. coli strains co-expressing CvFAP, with or without TrxA fusion. Protein loading was normalized based on cell density (OD600).

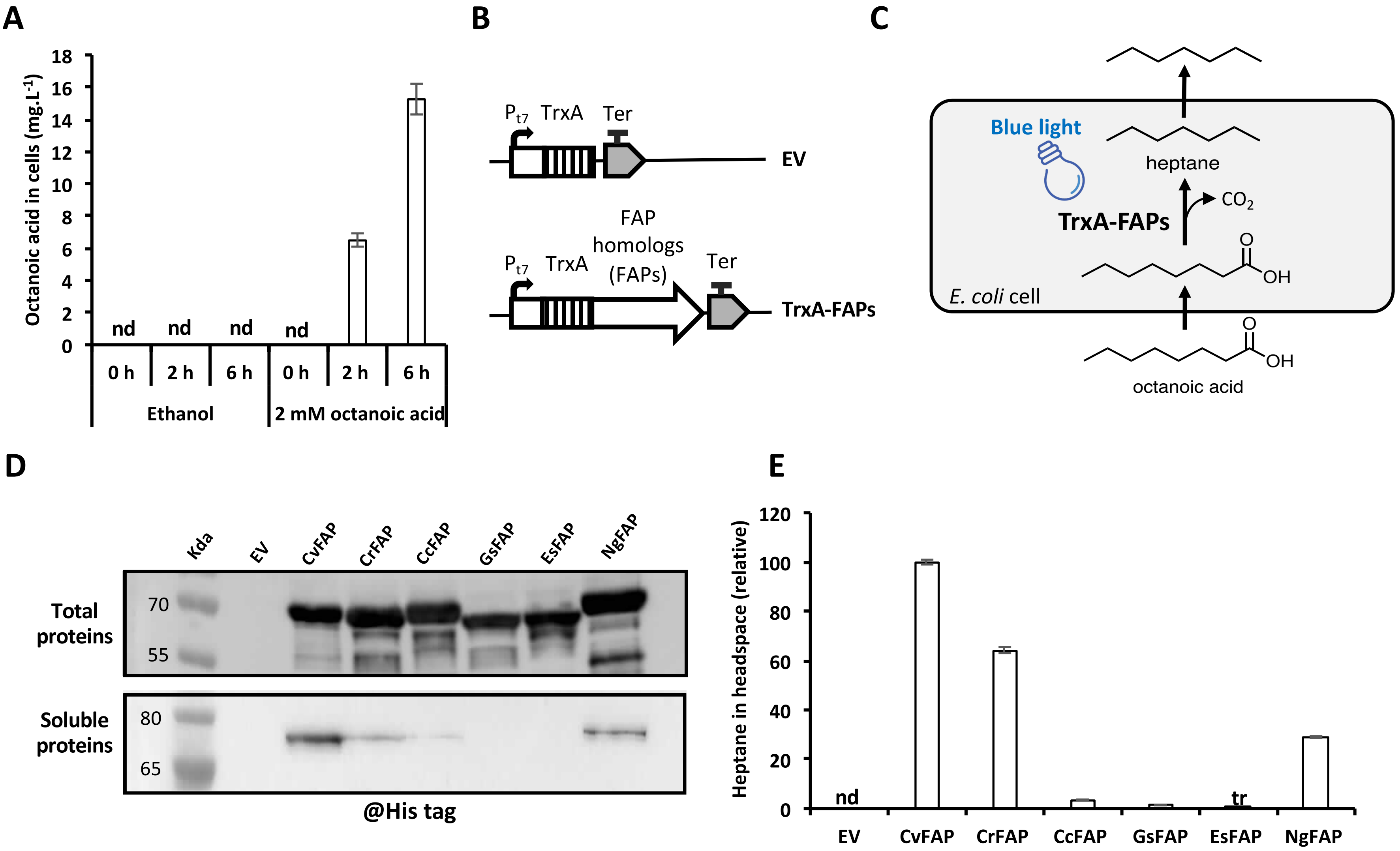


**Figure 2: Heptane Production in the Gas Phase of *E. coli* Cultures Expressing Various FAPs under Blue Light Illumination with Octanoic Acid Supplementation. A)** Time course of octanoic acid uptake by E. coli cells harbouring an empty vector (EV), treated with 2 mM octanoic acid (in ethanol) after 16 h of growth. Cells were washed twice prior to quantification. *E. coli* strains exposed only to ethanol served as controls. **B)** Schemes of the genetic constructs used in this study. **C)** Schematic representations of the metabolic pathways. **D)** Immunoblot detection of various FAP homologs in the total and soluble protein fractions from E. coli strains after a 16-hour culture period in the dark. Protein loading was normalized based on cell density (OD600). **E)** Relative quantification of heptane production in the gas phase of *E. coli* cultures expressing different FAP homologs, compared to control cultures with an EV. After growing in the dark for 16 hours, cultures were treated with 2 mM octanoic acid for 2 hours and then incubated under 300 µmol photons $m^{-2}$ $s^{-1}$ of blue light in sealed vials for 0.33 hours at 20°C. Error bars represent the standard error based on three biological replicates. The full immunoblot and protein gel are presented in Figure S3. 'nd'= not detected, 'tr'= trace.

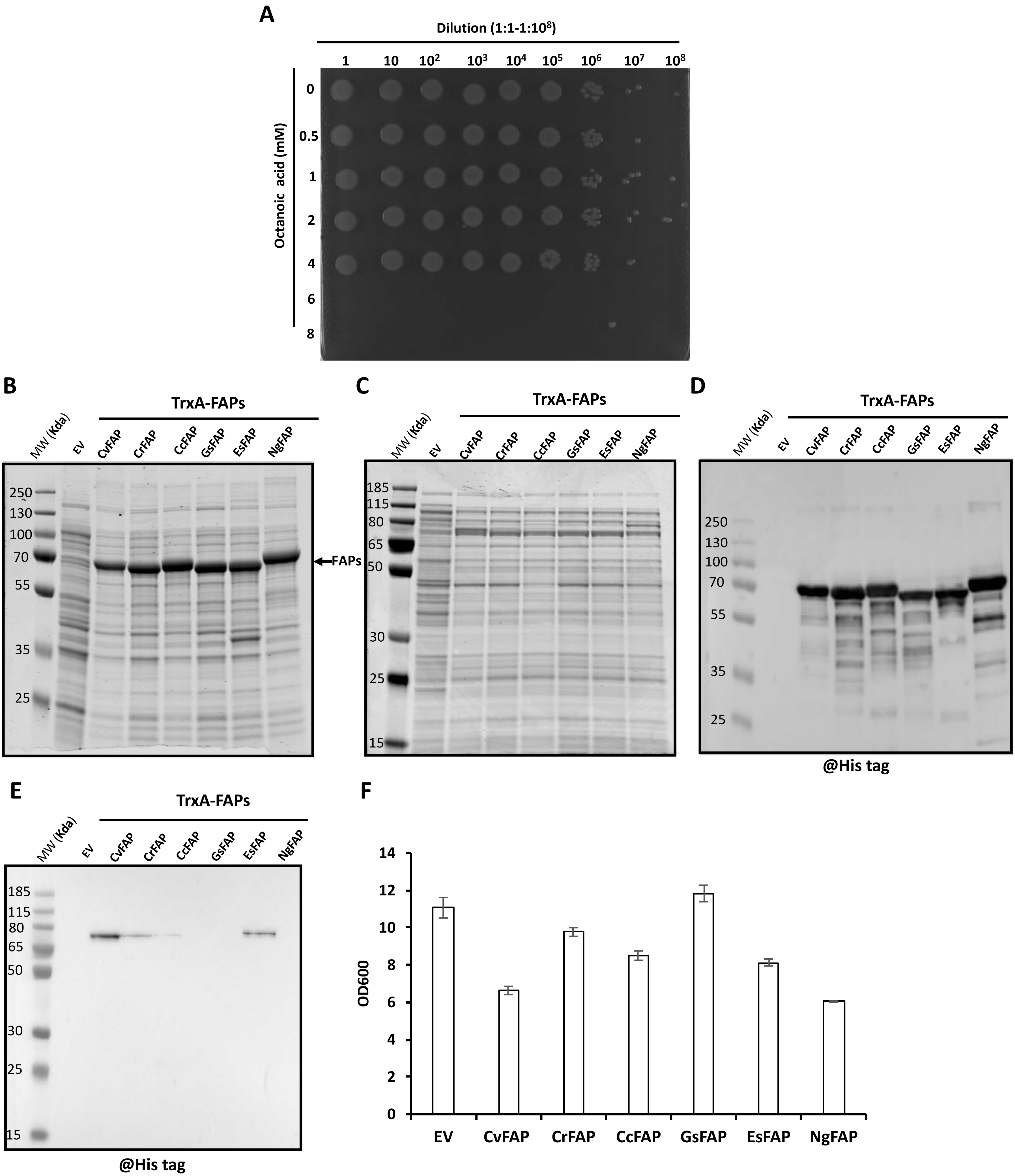


**Figure S3:** ***In Vivo* Synthesis of Heptane from Octanoic Acid by *E.coli* BL21(DE3) Expressing Various FAP Homologs. A)** Drop test of *E. coli* cells harbouring empty vector exposed to different concentrations of octanoic acid. After growing in the dark for 16 hours, cultures were treated with different concentrations of octanoic acid for 2 hours in dark and then incubated under 300 µmol photons $m^{-2} s^{-1}$ of blue light in sealed vials for 20 minutes at 20°C. SDS-PAGE analysis of total (**B**) and soluble (**C**) protein profiles from *E. coli* strains expressing different FAPs after a 16-hour culture period in the dark. Immunoblot detection of different FAPs in total protein extracts (**D**) or in the soluble protein fraction (**E**) *from E. coli* strains cultured for 16 hours in the dark. Protein loading was normalized based on cell density (OD600). **F)** Optical density (OD600) of *E. coli* cultures after a 16-hour cultivation period. Error bars represent the standard error based on three biological replicates. 'nd' = not detected.

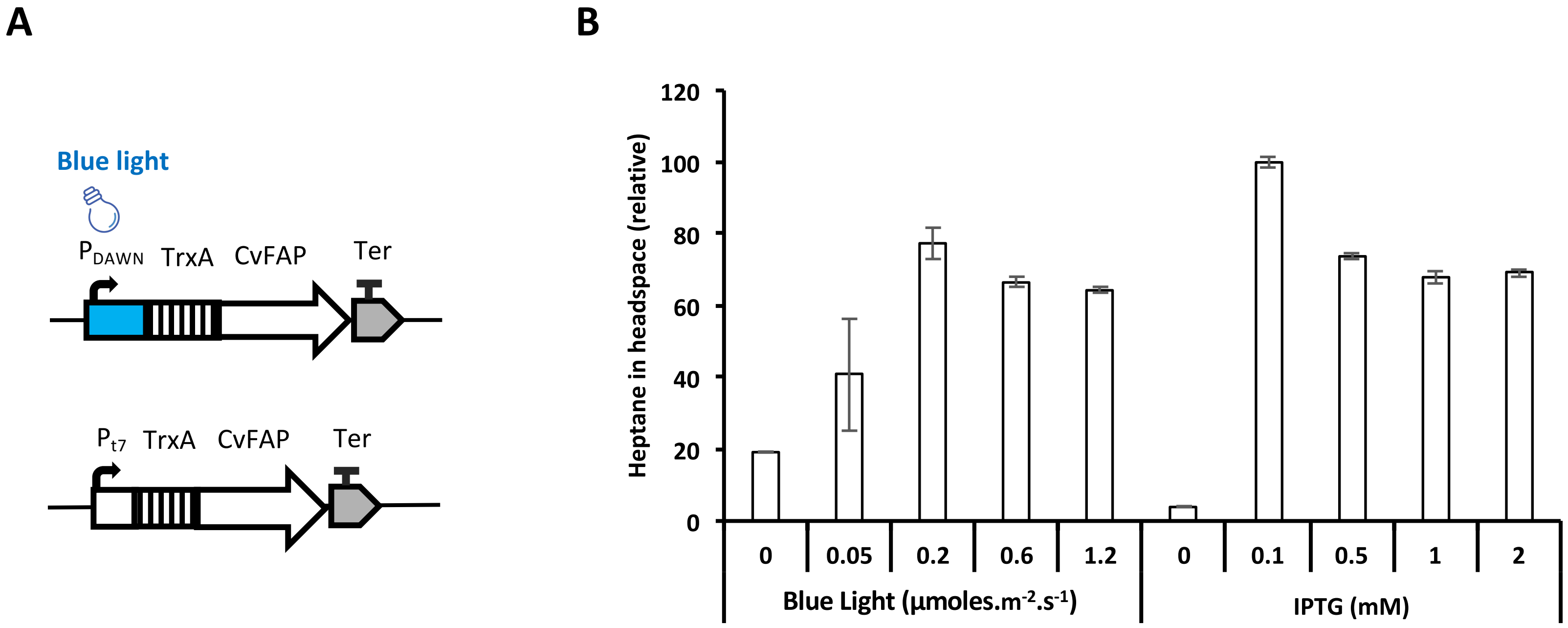


**Figure 3: Comparison of Hydrocarbon Productivity of BL21(DE3) E.coli strains expressing CvFAP under the control of Blue Light-Inducible and IPTG-Inducible Promoters. A)** Schemes of the genetic constructs used, featuring both blue light-inducible (PDAWN) and IPTG-inducible (Pt7) promoters. **B)** Relative quantification of heptane produced in the gas phase of *E. coli* cultures expressing *Cv*FAP under either PDAWN or Pt7. The strains expressed *Cv*FAP under the control of either PDAWN or Pt7 and were exposed to varying light intensities (0-1.2 µmoles.$m^{-2}.s^{-1}$) or different IPTG concentrations (0-2 mM), respectively during 16 h. Then, cultures were incubated with 2 mM octanoic acid for 2 hours in darkness, then exposed to 300 µmol photons $m^{-2}s^{-1}$ of blue light in sealed vials for a 0.33 hours period at 20°C. Error bars represent the standard error based on five biological replicates.

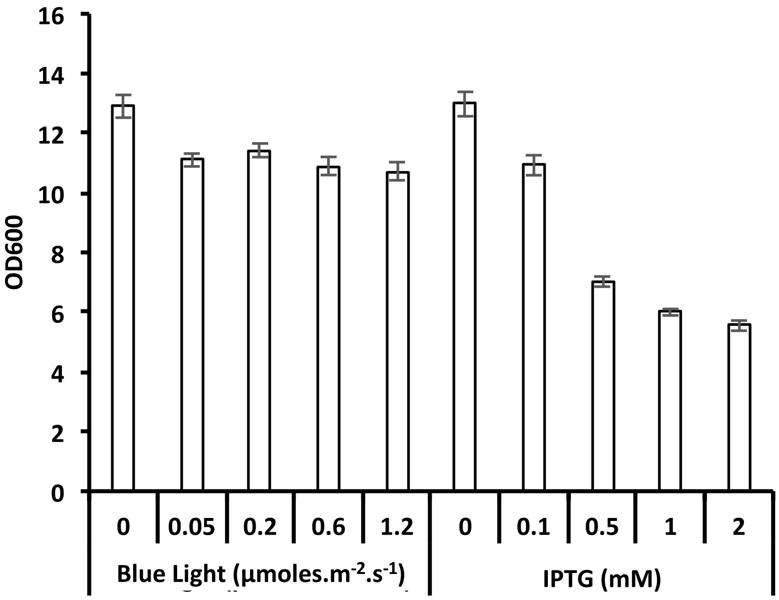


**Figure S4: Growth of *E. coli* cells expressing CvFAP under either the blue light-inducible promoter or the IPTG-inducible promoter, after a 16-hour culture period. The cells were exposed to light intensities ranging from 0 to 1.2 µmol photons.m$^{-2}$.s$^{-1}$ or IPTG concentrations from 0 to 2 mM.** Error bars represent the standard deviation based on three biological replicates.

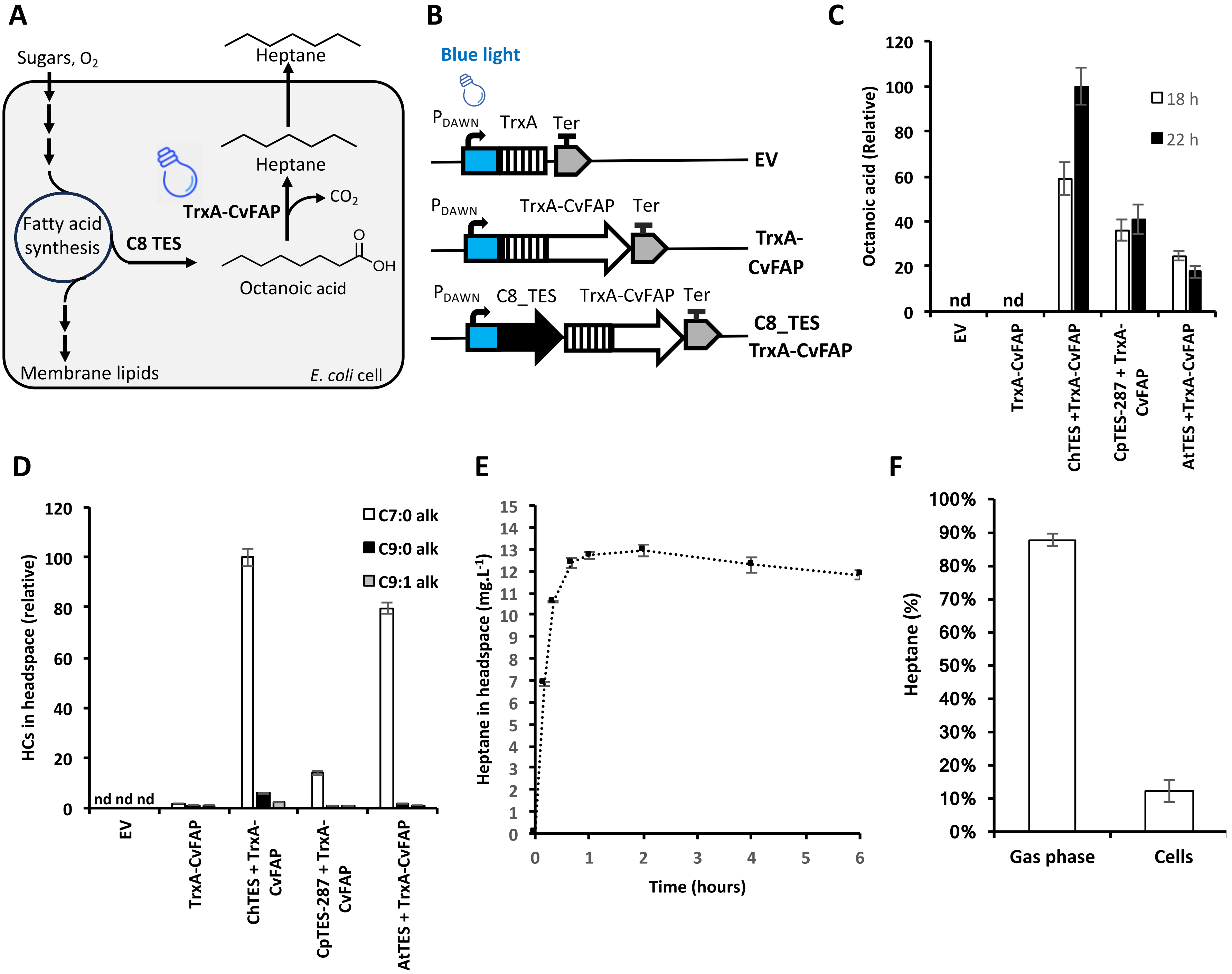


**Figure 4**: **Co-expression of Various Octanoyl-ACP Substrate-Specific Thioesterases (C8TES) and CvFAP Genes Under Blue-Light Inducible Promoter in *E. coli* BL21(DE3). A)** Schematic representation of the metabolic pathways involved in heptane formation. **B)** Scheme of the genetic constructs. **C)** Octanoic acid content of *E. coli* cells co-expressing *Cv*FAP and different His-tagged C8TES before (18 h) and after (24 h) the illumination of the cultures. **D)** Volatile hydrocarbon levels measured in the headspace produced by *E. coli* cells co-expressing ChTES and CvFAP genes, following 4 hours of incubation under 300 µmol photons.m$^{-2}$.s$^{-1}$. **E)** Kinetics of heptane production in the headspace of *E. coli* strains co-expressing His-tagged *Ch*TES and *Cv*FAP after illumination at 300 µmol photons.m$^{-2}$.s$^{-1}$ . **F)** Proportion of heptane present in the headspace and inside cells produced by *E. coli* cells co-expressing the His-tagged *Ch*TES and *Cv*FAP genes after 4 h of incubation at 300 µmol photons.m$^{-2}$.s$^{-1}$. Error bars represent the standard error based on three biological replicates. 'nd' = non detected.

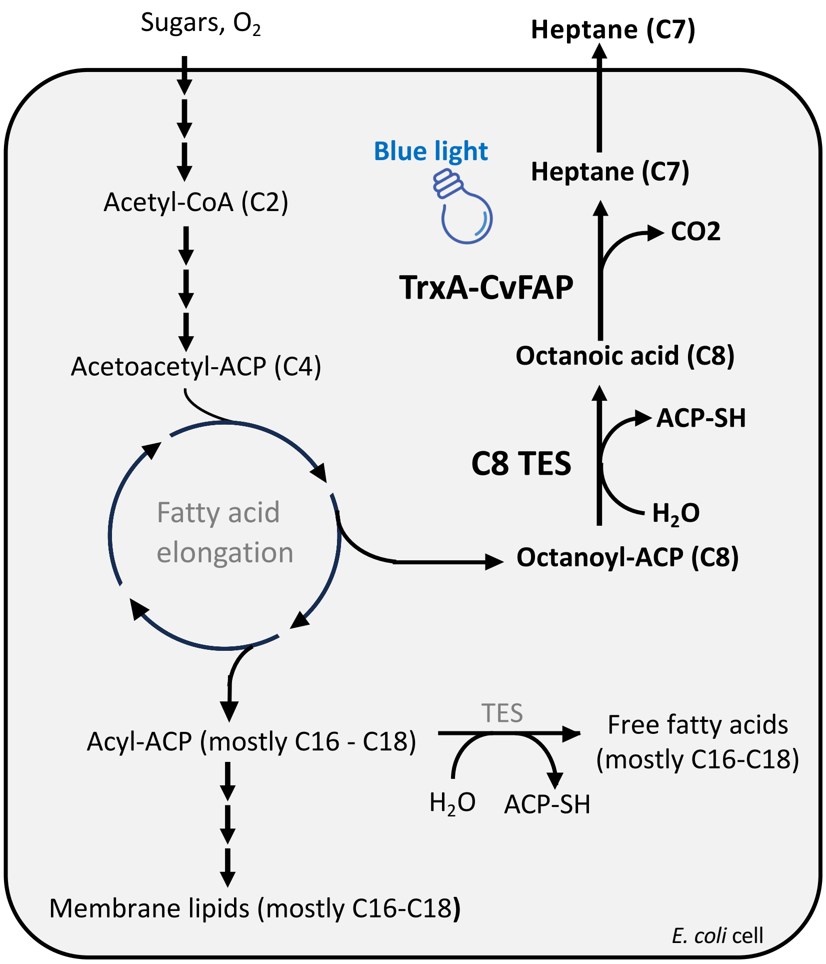


**Figure S5: Metabolic Pathway Map in *E. coli* for the Synthesis of Octanoic Acid and Heptane.** This figure outlines the enzymatic reactions within *E. coli* fatty acid metabolism relevant to octanoic acid and heptane production. Bold black pathways represent the heterologous pathways introduced into E. coli to direct fatty acid biosynthesis towards heptane production. **CvFAP:** Chlorella variabilis fatty acid photodecarboxylase, **C8_TES:** Octanoyl-ACP specific thioesterase, **TES:** Endogenous *E. coli* thioesterases, **CoA:** Coenzyme A
**ACP:** Acyl-Carrier Protein

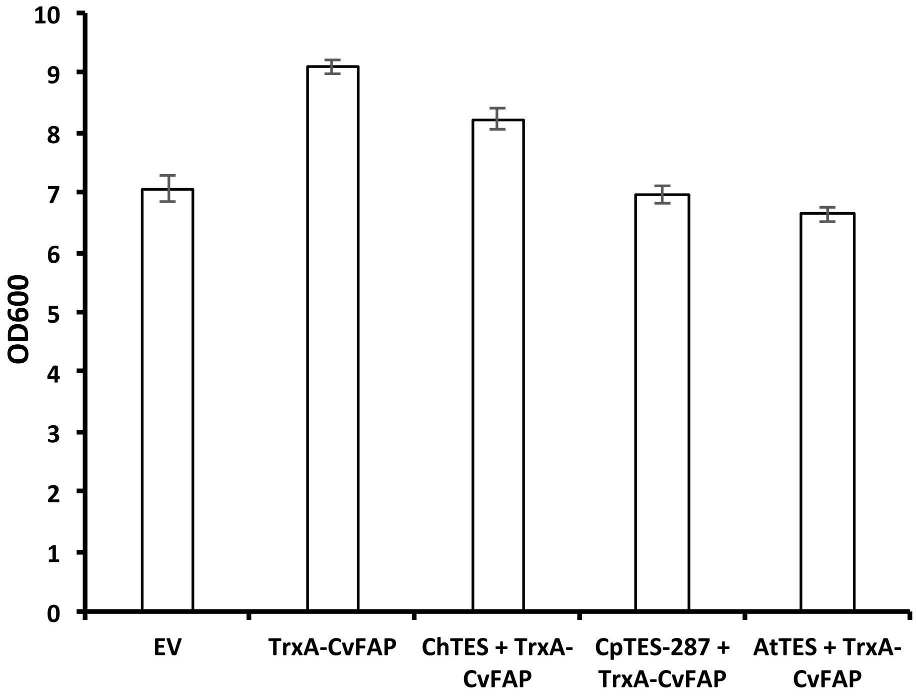


**Figure S6: Optical Density Measurements of *E. coli* Strains Co-expressing Various His-tagged C8_TES and CvFAP.** This figure displays the optical density of E. coli strains after a 18-hour cultivation period and exposed to 0.2 µmol photons.$m^{-2}.s^{-1}$. Error bars represent the standard error based on three biological replicates.

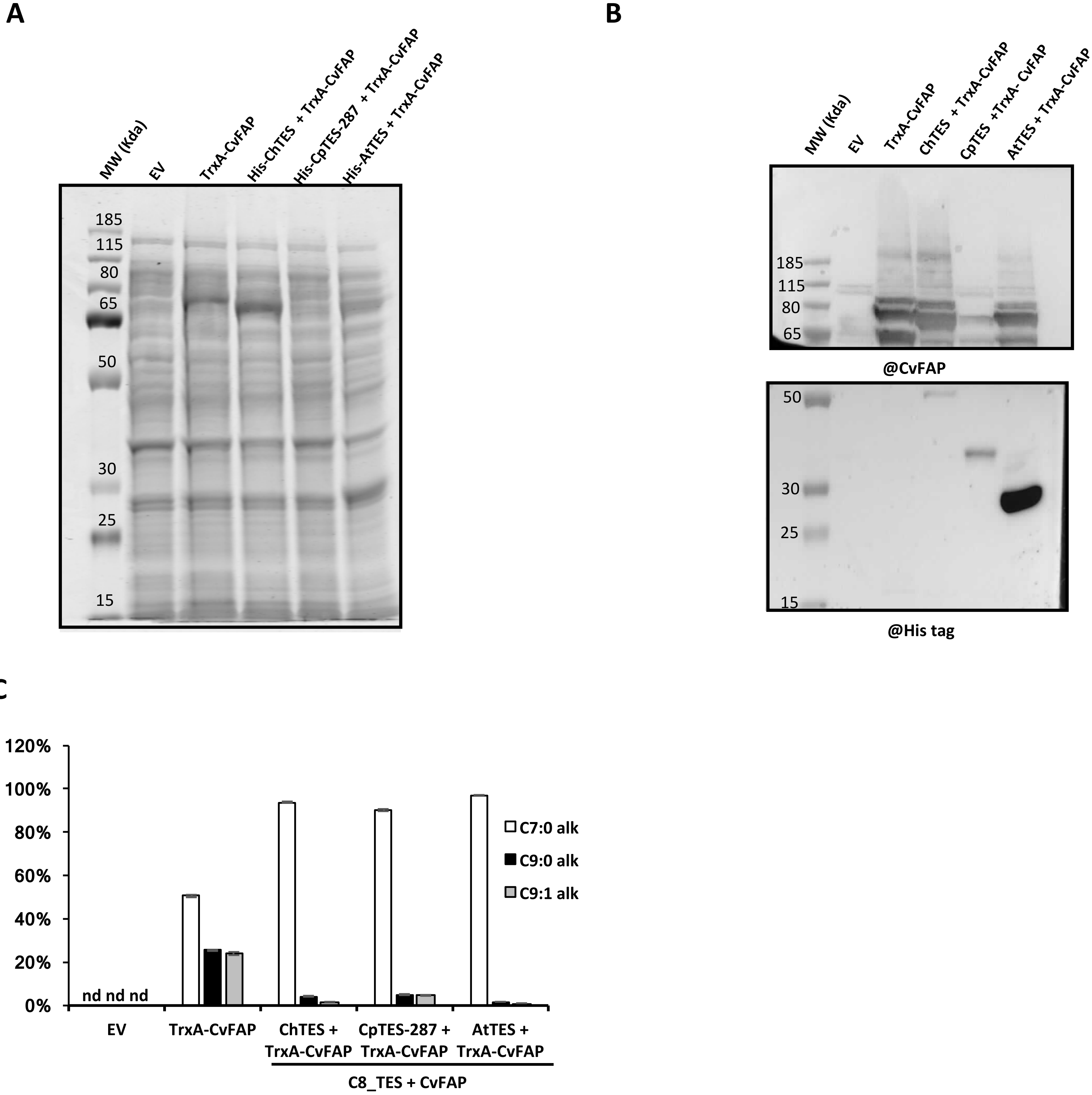


**Figure S7: Production of Volatile Hydrocarbons by *E. coli* BL21(DE3) Expressing Different Octanoyl-ACP Specific Thioesterases (C8_TES) and CvFAP Under Blue-Light Inducible Promoter. A)** SDS-PAGE analysis of the total protein profile from E. coli strains harvested after an 18-hour culture period. **B)** Immunoblot analysis of total proteins from *E. coli* strains after 18 hours of culture: Upper panel shows detection of CvFAP, lower panel shows detection of His-tagged C8_TES thioesterases. **C)** Composition of volatile hydrocarbons produced in the headspace of sealed vials by *E. coli* strains co-expressing different His-tagged C8_TES and CvFAP, measured after 4 hours of illumination at 300 µmol photons.$m^{-2}$.$s^{-1}$. Error bars represent the standard error based on three biological replicates. nd = not detected.

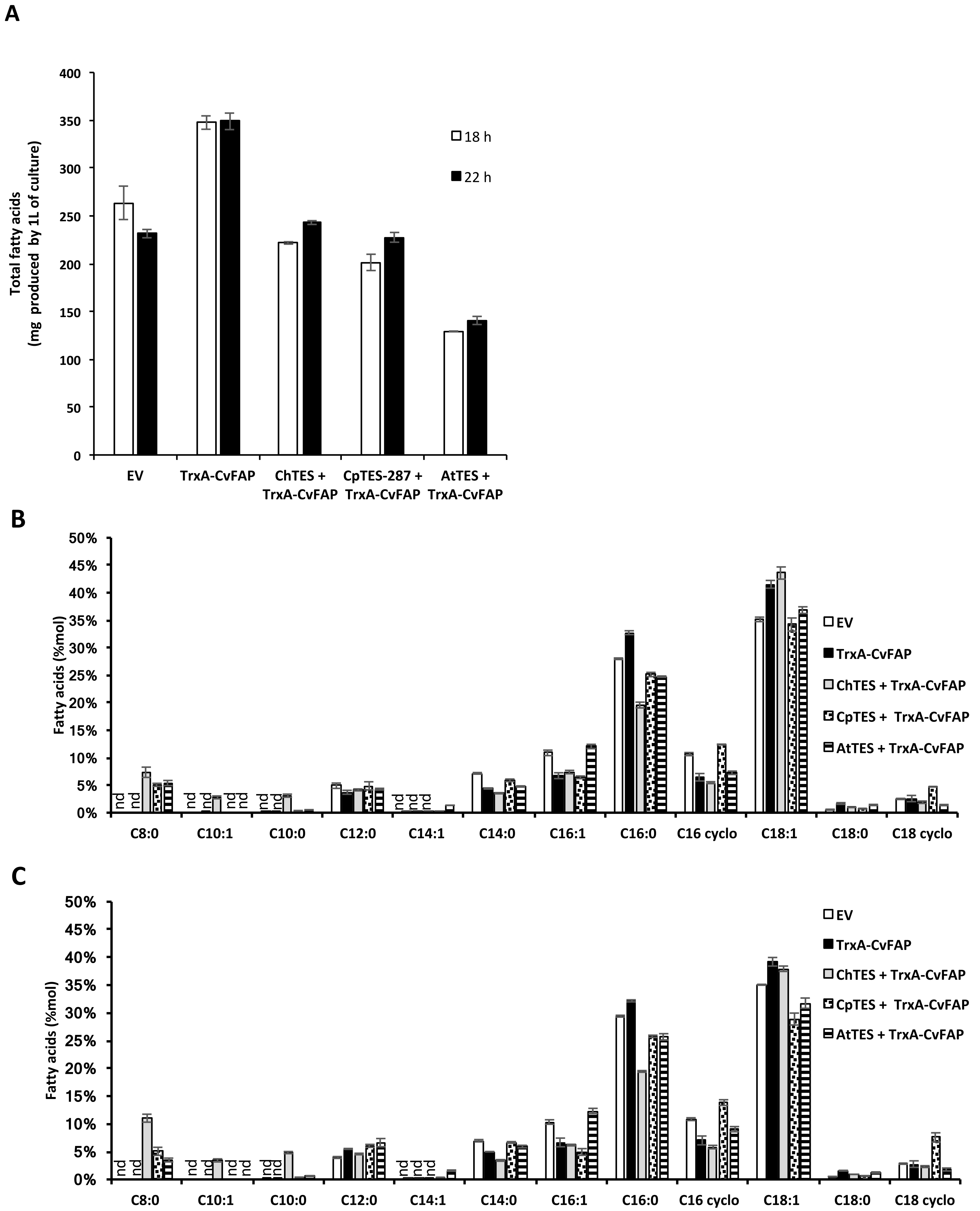


**Figure S8: Fatty Acid Profiles in *E. Coli* BL21(DE3) Co-expressing Different Octanoyl-ACP Substrate-Specific Thioesterases (C8_TES) and CvFAP Under a Blue-Light Inducible Promoter. A)** Total fatty acid content in *E. coli* cells before and after 4 hours of incubation at 300 µmol photons $m^{-2} s^{-1}$. Control strains carried an empty vector (EV). Fatty acid composition of *E. coli* cells, measured before **(B)** and after **(C)** the 4-hour incubation period at 300 µmol photons $m^{-2} s^{-1}$. Comparison is made against the EV control. Error bars represent the standard error from three biological replicates. 'nd' = not detected.

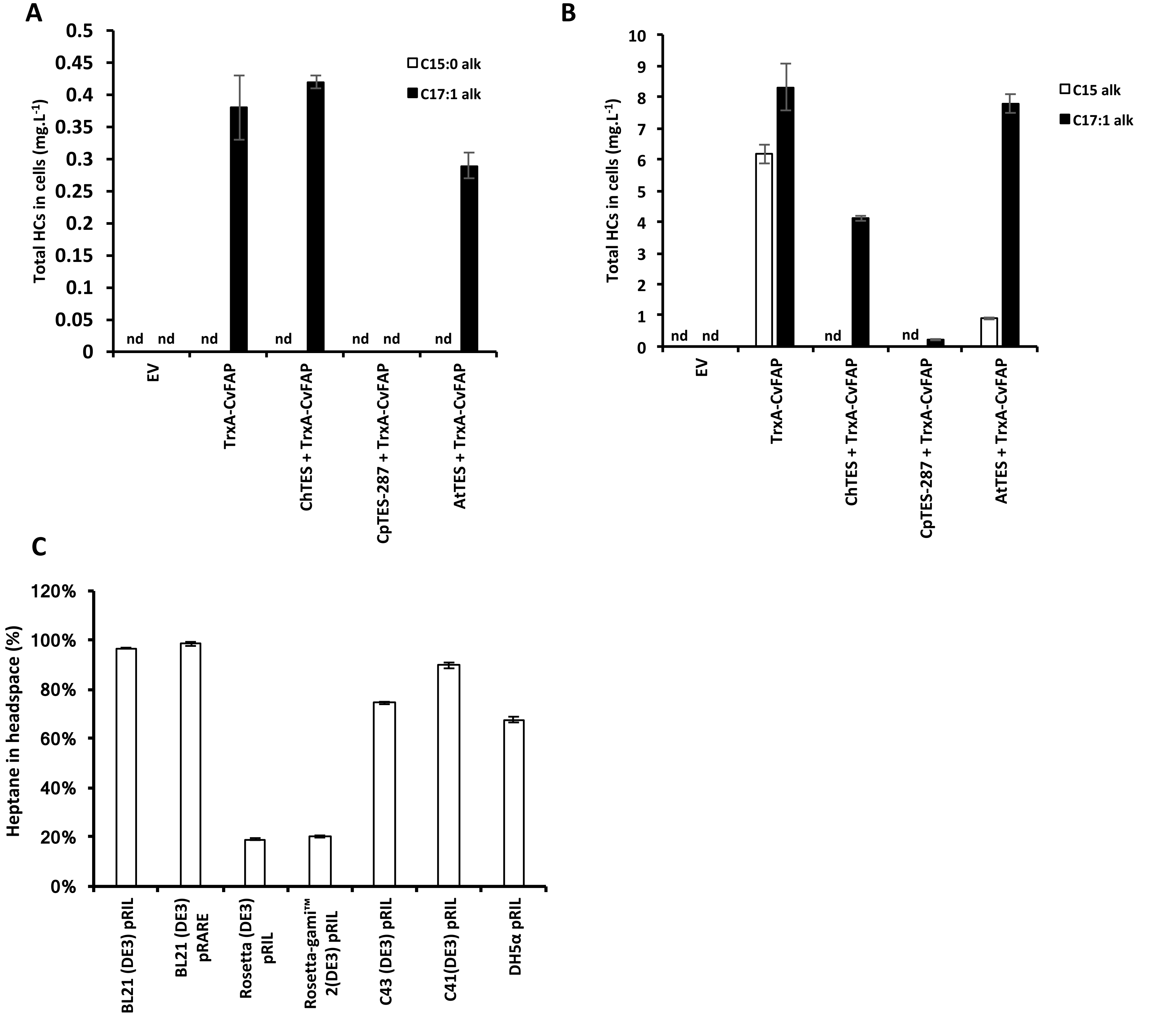


**Figure S9: Hydrocarbon Production by *E. coli* BL21(DE3) Co-expressing Octanoyl-ACP Substrate-Specific Thioesterases (C8_TES) and CvFAP Under a Blue-Light Inducible Promoter.** Hydrocarbon content inside *E. coli* cells measured before **(A)** and after **(B)** a 4-hour incubation period at 300 µmol. $m^{-2}·s^{-1}$. **B)** Composition of hydrocarbon content inside *E. coli* cells, assessed before and after the 4-hour incubation period at 300 µmol. $m^{-2}·s^{-1}$. **C)** Relative production of heptane in the headspace by different *E. coli* strains co-expressing ChTES and CvFAP, compared after 4 hours of incubation at 300 µmol. $m^{-2}·s^{-1}$. All experimental data were compared with an empty vector (EV) as a control. Error bars represent the standard error from three biological replicates. 'nd'= not detected.

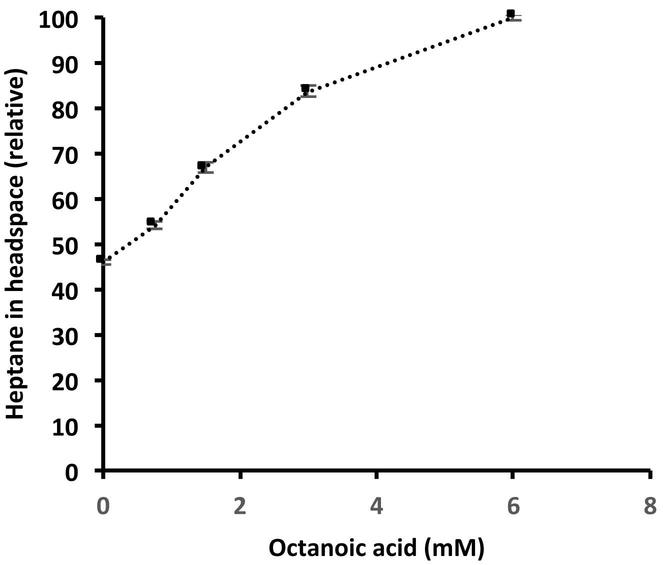


**<u>Figure 5</u>: Detection of heptane in *E. coli* strains co-expressing ChTES and CvFAP genes, fed with varying concentrations of octanoic acid and incubated under 300 µmol photons.$m^{-2}.s^{-1}$ of blue light in sealed vials for 4 hours at 20°C**. Error bars represent the standard error based on three biological replicates.

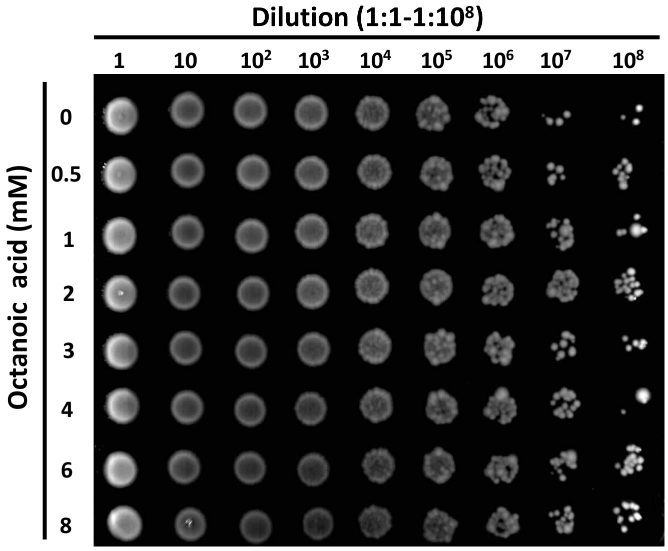


**<u>Figure S10</u>: Drop test of *E. coli* cells strains co-expressing ChTES and CvFAP exposed to different concentrations of octanoic acid (0-8 mM).** After growing in the cells for 16 hours incubated under 0.2 µmol photons $m^{-2}$ $s^{-1}$ of blue light, cultures were treated with different concentrations of octanoic acid for 2 hours in dark and then incubated under 300 µmol photons $m^{-2}$ $s^{-1}$ of blue light in sealed vials for 4 hours at 20°C.

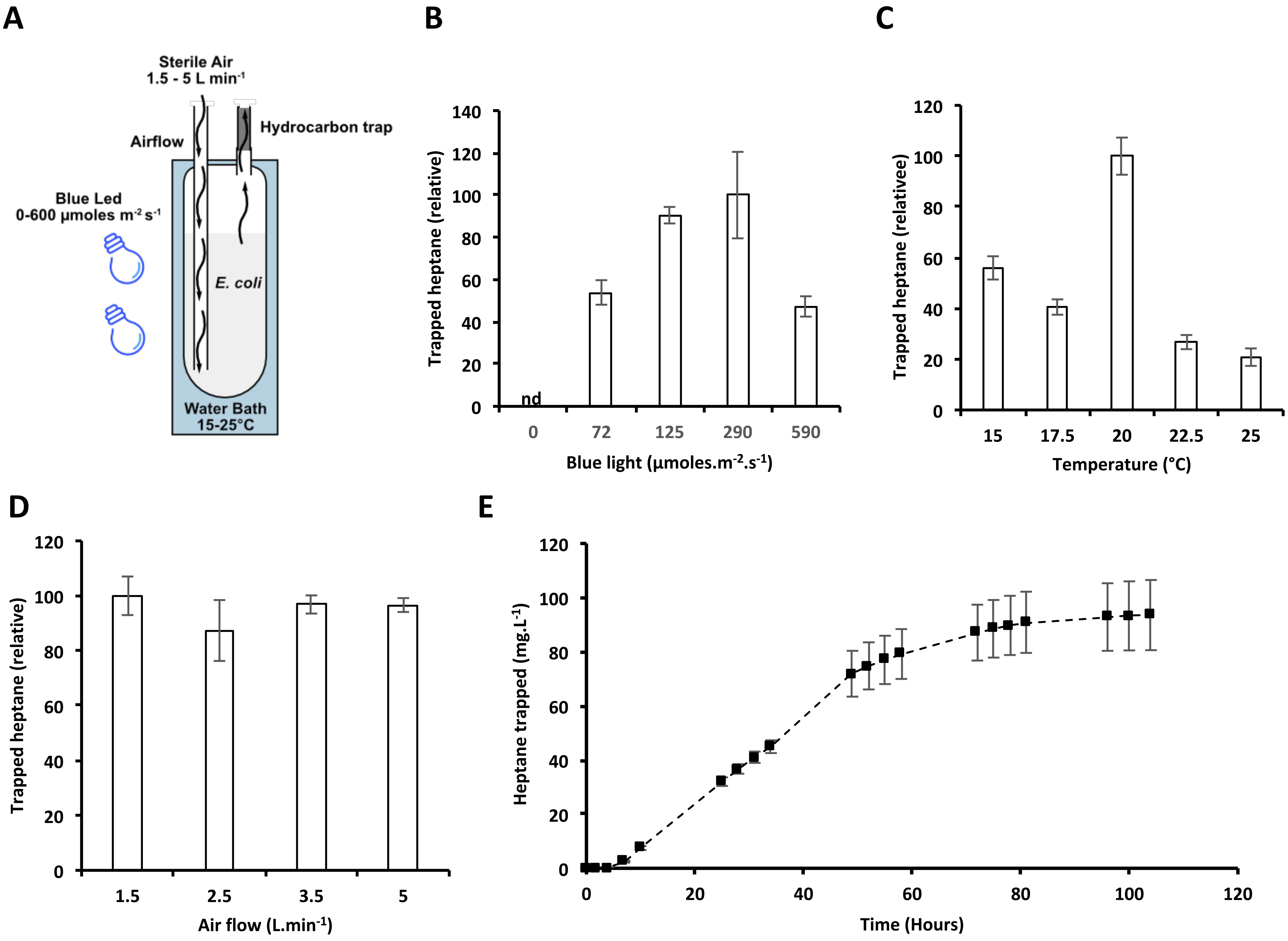


**Figure 6: Optimization of Continuous Photoproduction of Heptane in *E.coli* BL21(DE3) Co-expressing *Ch*TES and *Cv*FAP in a 100 ml Photobioreactor. A)** Schematic of the photobioreactor system utilized for heptane production. This system includes adjustable settings for light intensity, temperature, and airflow, and is equipped desorption liners packed with Tenax TA™, which retains volatile compounds including medium-chain hydrocarbons like heptane. **B)** Relative heptane production from *E. coli* strains cultured for 18 hours at 20°C, with an airflow of 5 L/min, across various blue light intensities (0-590 µmol photons.$m^{-2}.s^{-1}$). **C)** Relative heptane production from strains cultured under 290 µmol photons.$m^{-2}.s^{-1}$ of blue light exposure for 18 hours at different temperatures. **D)** Relative heptane production from strains cultured under similar conditions with varying airflow rates in the photobioreactor. **E)** Time course of cumulative heptane production in *E. coli* grown under continuous blue light illumination (290 µmol photons.$m^{-2}.s^{-1}$) for 104 hours at 20°C and an airflow rate of 1.5 L/min. Error bars represent the standard error from multiple biological replicates (parts B and E n=6, part C n=7, part D n=4). nd = not detected.

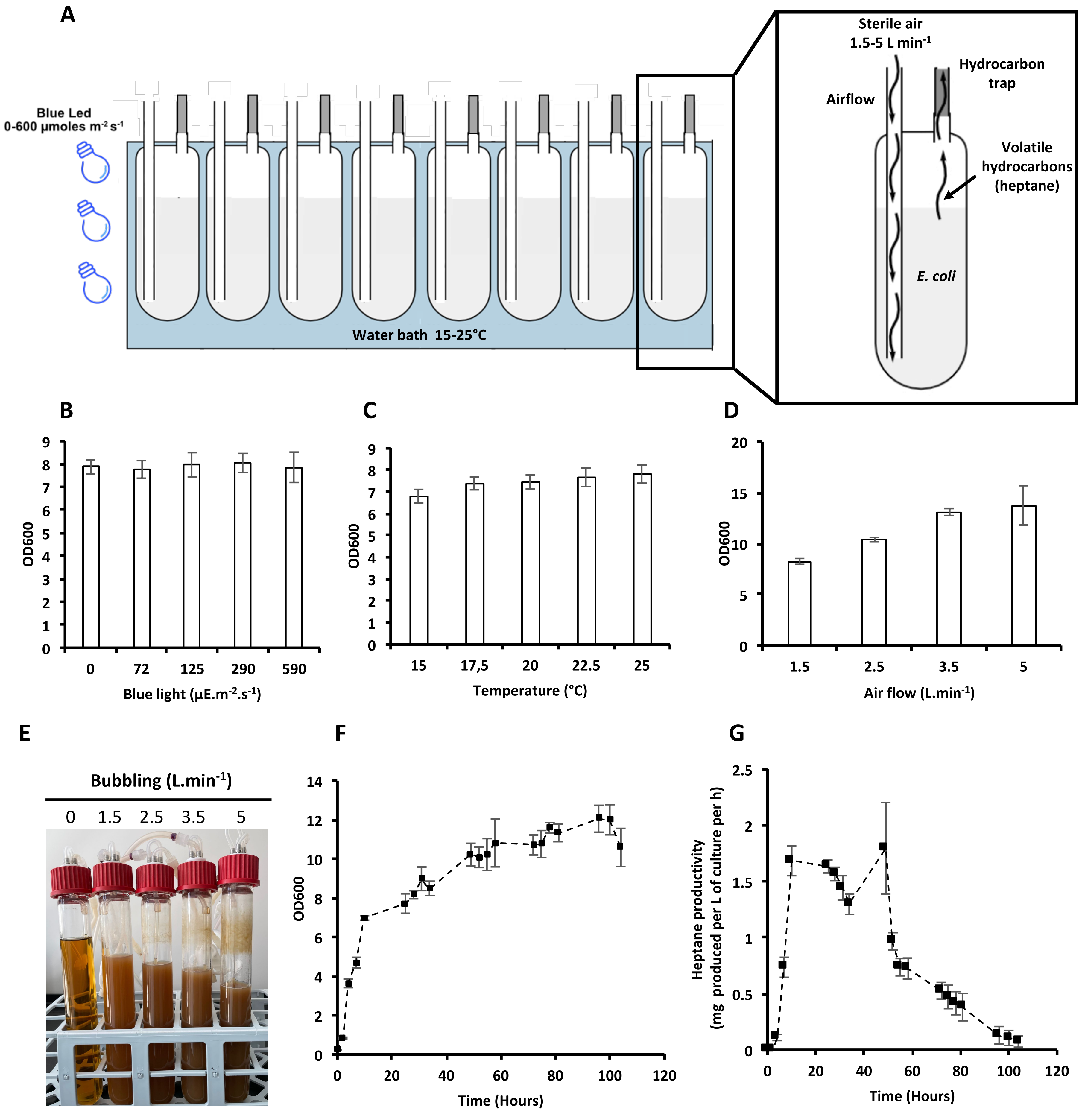


**<u>Figure S11</u>: Continuous Photoproduction of Volatile Heptane in *E.coli* BL21(DE3) Expressing CvFAP and ChTES in a 100 ml Photobioreactor. A)** Schematic representation of the 100 ml photobioreactor system used for heptane production. The system modulates various culture parameters including light intensity, temperature, and airflow, and includes a thermodesorption liner packed with Tenax TA™ for retaining volatile compounds like medium-chain hydrocarbons. **B)** Optical density of *E. coli* strains cultured for 18 hours at 20°C with an airflow of 5 $L.min^{-1}$, exposed to varying blue light intensities (0-590 $\mu mol.m^{-2}.s^{-1}$). **C)** Optical density of strains cultured for 18 hours under 290 $\mu mol.m^{-2}.s^{-1}$ blue light, with airflow of 5 $L.min^{-1}$, at different temperatures. **D)** Optical density of strains cultured for 18 hours at 20°C under 290 $\mu mol.m^{-2}.s^{-1}$ blue light, with varying airflow rates. **E)** Photograph illustrating the evaporation of E. coli cultures over 3 days under conditions detailed in (D). **F)** Time course of optical density in *E. coli* grown under constant blue light illumination (290 $\mu mol.m^{-2}.s^{-1}$) for 104 hours at 20°C with an airflow of 1.5 $L.min^{-1}$. **G)** Heptane productivity calculated at each time point from the experiment shown in Figure 5F. Error bars represent the standard deviation (parts B, C, D, and E) or standard error (parts F and G) from multiple biological replicates, part (B and F) n=6, part (C) n=7, part (D) n=4

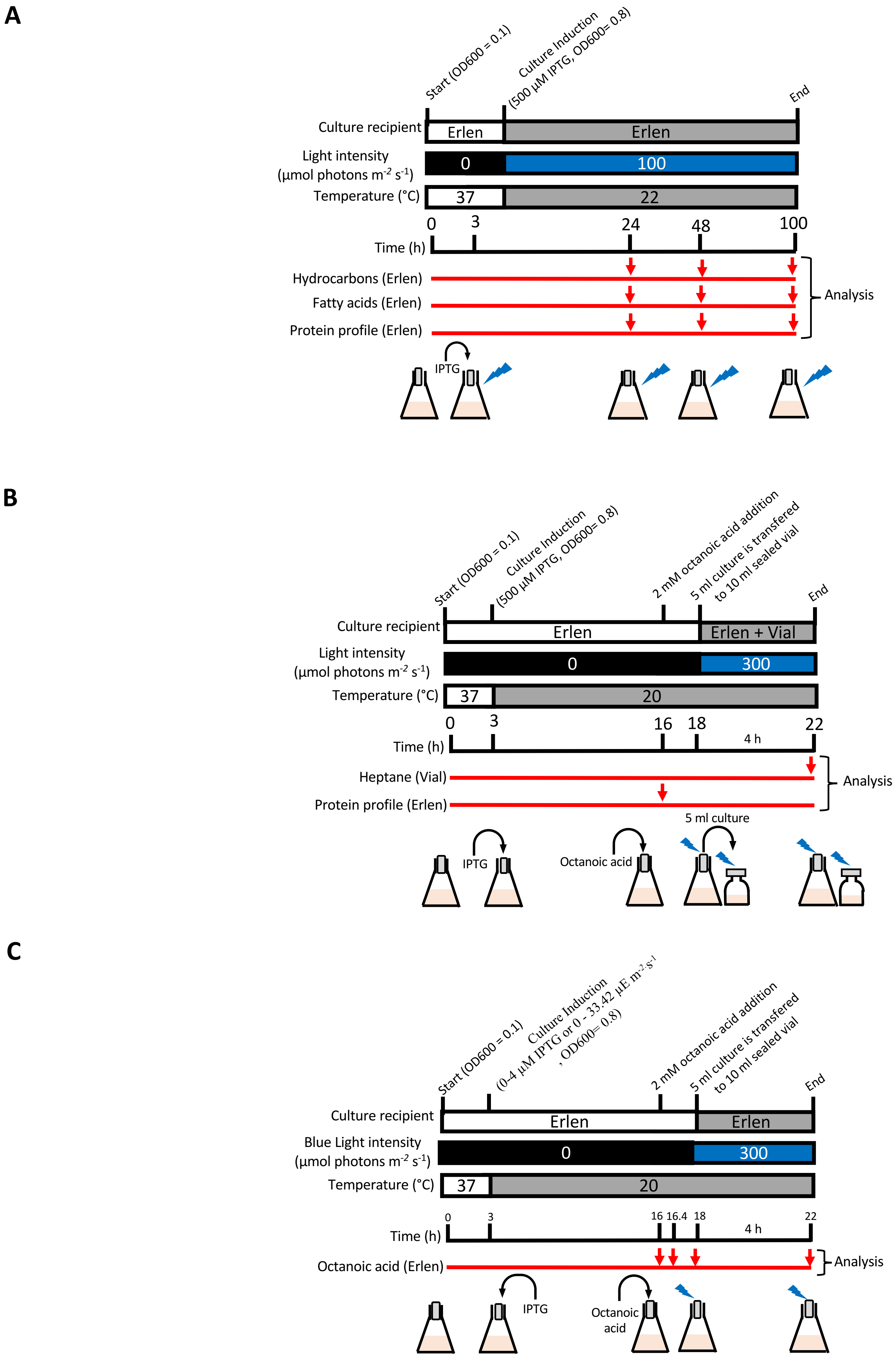


**Figure S12: Experimental design. A)** Experimental design for the experiment proposed in **Figure 1** (A), in **Figure 2A, 2C and 2D** (B) and in **Figure 2B** (C).

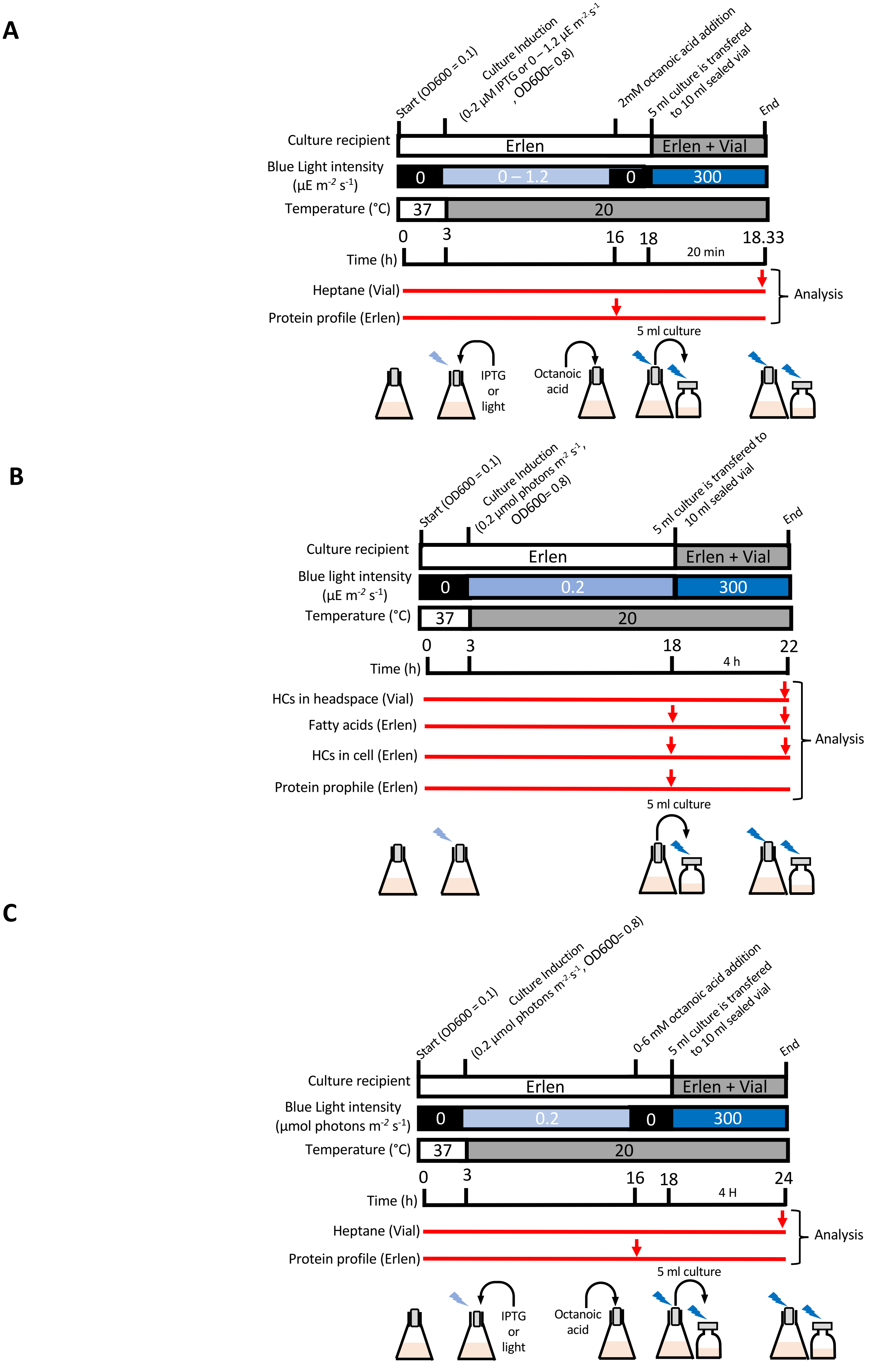


**Figure S13: Experimental design A)** Experimental design for the experiment proposed in **Figure 3**. (**A**), in **Figure 4** (except **Figure 4F**) (**B**) and **Figure 4F** (**C**)

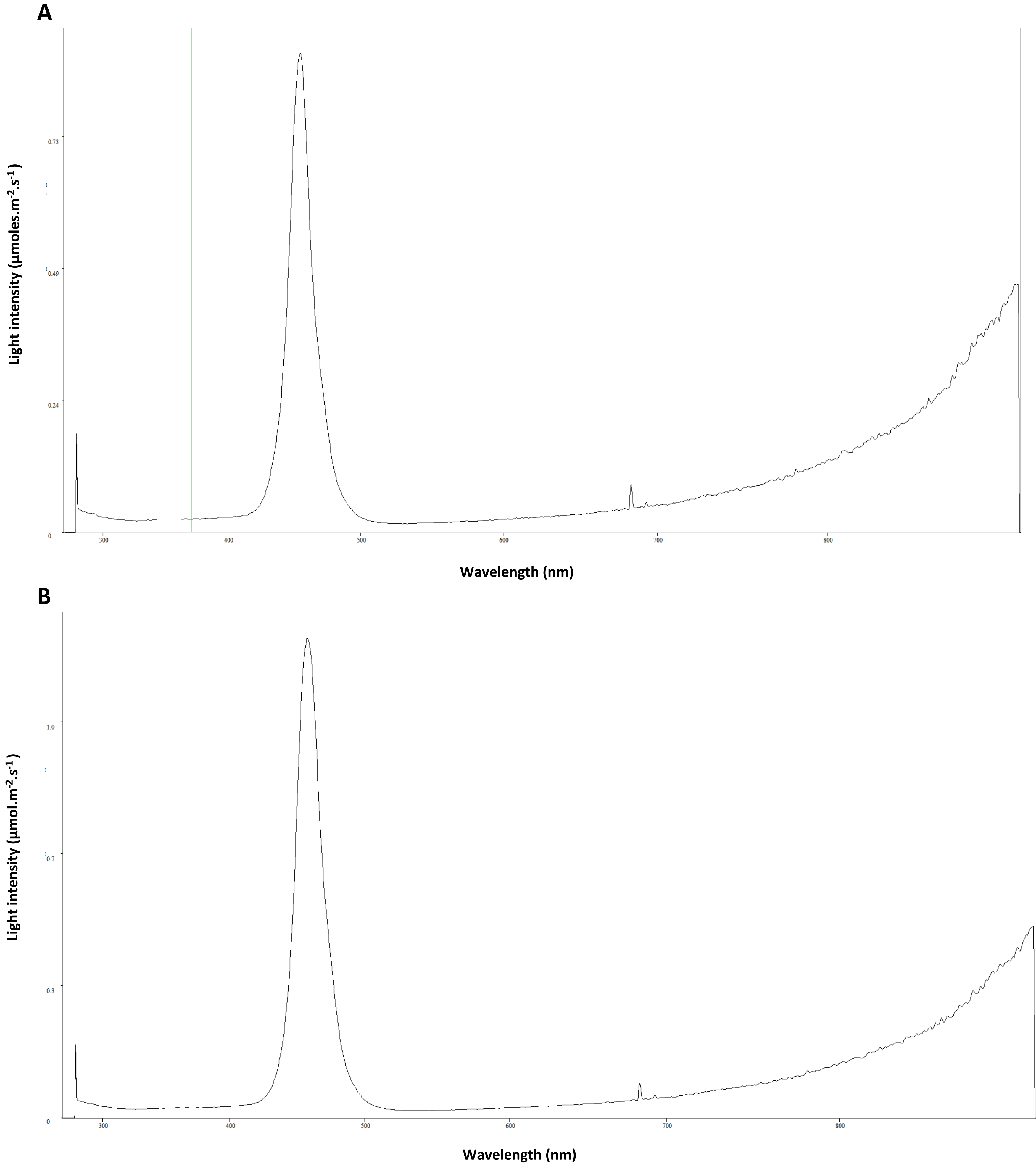


<u>Figure S14</u>**: The different LED blue light spectra used in this work. A)** Spectra of the blue light emitted by the LED panel used for the experiments presented in **Figure 2, 3, 4 .** **B)** Spectra of the blue light emitted by the LED panel used for the experiments presented in **Figure 5 and 6.**

| Protein | Full name | Species | Aminoacid sequence (N-terminal to C-terminal) in FASTA format |
|---|---|---|---|
| **CvFAP** | Fatty acid photodecarboxylase | *Chlorella variabilis* | >CvFAP<br>MGHHHHHHSSGVDLGTENLYFQSMASAVEDIRKVLSDSSSPVAGQKYDYILVGGGTAACVLANRLSADGSKRVLVLEAGPDNTSRDVKIPAAITRLFRSP<br>LDWNLFSELQEQLAERQIYMARGRLLGGSSATNATLYHRGAAGDYDAWGVEGWSSEDVLSWFVQAETNADFGPGAYHGSGGPMRVENPRYTNKQL<br>HTAFFKAAEEVGLTPNSDFNDWSHDHAGYGTFQVMQDKGTRADMYRQYLKPVLGRRNLQVLTGAAVTKVNIDQAAGKAQALGVEFSTDGPTGERLS<br>AELAPGGEVIMCAGAVHTPFLLKHSGVGPSAELKEFGIPVVSNLAGVGQNLQDQPACLTAAPVKEKYDGIAISDHIYNEKGQIRKRAIASYLLGGRGGLTST<br>GCDRGAFVRTAGQALPDLQVRFVPGMALDPDGVSTYVRFAKFQSQGLKWPSGITMQLIACRPQSTGSVGLKSADPFAPPKLSPGYLTDKDGADLATLR<br>KGIHWARDVARSSALSEYLDGELFPGSGVVSDDQIDEYIRRSIHSSNAITGTCKMGNAGDSSSVVDNQLRVHGVEGLRVVDASVVPKIPGGQTGAPVVM<br>IAERAAALLTGKATIGASAAAPATVAA |
| **TrxA-CvFAP** | | *Chlorella variabilis* | >TrxA-CvFAP<br>MGHHHHHHSDKIIHLTDDSFDTDVLKADGAILVDFWAEWCGPCKMIAPILDEIADEYQGKLTVAKLNIDQNPGTAPKYGIRGIPTLLLFKNGEVAATKVG<br>ALSKGQLKEFLDANLAGSGSGVSGVDLGTENLYFQSASAVEDIRKVLSDSSSPVAGQKYDYILVGGGTAACVLANRLSADGSKRVLVLEAGPDNTSRDVKI<br>PAAITRLFRSPLDWNLFSELQEQLAERQIYMARGRLLGGSSATNATLYHRGAAGDYDAWGVEGWSSEDVLSWFVQAETNADFGPGAYHGSGGPMRV<br>ENPRYTNKQLHTAFFKAAEEVGLTPNSDFNDWSHDHAGYGTFQVMQDKGTRADMYRQYLKPVLGRRNLQVLTGAAVTKVNIDQAAGKAQALGVEFS<br>TDGPTGERLSAELAPGGEVIMCAGAVHTPFLLKHSGVGPSAELKEFGIPVVSNLAGVGQNLQDQPACLTAAPVKEKYDGIAISDHIYNEKGQIRKRAIASYL<br>LGGRGGLTSTGCDRGAFVRTAGQALPDLQVRFVPGMALDPDGVSTYVRFAKFQSQGLKWPSGITMQLIACRPQSTGSVGLKSADPFAPPKLSPGYLTDK<br>DGADLATLRKGIHWARDVARSSALSEYLDGELFPGSGVVSDDQIDEYIRRSIHSSNAITGTCKMGNAGDSSSVVDNQLRVHGVEGLRVVDASVVPKIPGG<br>QTGAPVVMIAERAAALLTGKATIGASAAAPATVAA |
| **TrxA-CrFAP** | | *Chlamydomonas reinhardtii* | >TrxA-CrFAP<br>MGHHHHHHSDKIIHLTDDSFDTDVLKADGAILVDFWAEWCGPCKMIAPILDEIADEYQGKLTVAKLNIDQNPGTAPKYGIRGIPTLLLFKNGEVAATKVG<br>ALSKGQLKEFLDANLAGSGSGVSGVDLGTENLYFQSMEVAVPAGITRLFAHPVMDWGMSSLTQKQLVAREIYLARGRMLGGSSGSNATLYHRGSAADY<br>DAWGLEGWSSKDVLDWFVKAECYADGPKPYHGTGGSMNTEQPRYENVLHDEFFKAAAATGLPANPDFNDWSHPQDGFGEFQVSQKKGQRADTYR<br>TYLKPAMARGNLKVVIGARATKVNIEKGSSGARTTGVEYAMQQFGDRFTAELAPGGEVLMCSGAVHTPHLLMLSGVGPAATLKEHGIDVVSDLSGVGQ<br>NLQDHPAAVLAARAKPEFEKLSVTSEVYDDKCNIKLGAVAQYLFQRRGPLATTGCDHGAFVRTSSSLSQPDLQMRFVPGCALDPDGVKSYIVFGELKKQG<br>RAWPGGITLQLLAIRAKSKGSIGLKAADPFINPAININYFSDPADLATLVNAVKMARKIAAQEPLKKYLQEETFPGERASSDKDLEEYIRRTVHSGNALVGTA<br>AMGASPAAGAVVSSADLKVFGVEGLRVVDASVLPRIPGGQTGAATVMVAERAAALLRGQATIAPSRQPVAV |
| **TrxA-CcFAP** | | *Chondrus crispus* | >TrxA-CcFAP<br>MGHHHHHHSDKIIHLTDDSFDTDVLKADGAILVDFWAEWCGPCKMIAPILDEIADEYQGKLTVAKLNIDQNPGTAPKYGIRGIPTLLLFKNGEVAATKVG<br>ALSKGQLKEFLDANLAGSGSGVSGVDLGTENLYFQSSSEAATTYDYIIVGGGAAGCVLANRLTEDPSTRVLLLEAGKPDDSFYLHVPLGFPYLLGSPNDWAF<br>VTEPEPNLANRRLYFPRGKVLGGSHAISVMLYHRGHPADYTAWAESAPGWAPQDVLPYFLKSESQQSAVPNQDAHGYEGPLAVSDLARLNPMSKAFIK<br>AAHNAAGLNHNPDFNDWATGQDGVGPFQVTQRDGSRESPATSYLRAAKGRRNLTVMTGAVVERILFENPAGSSTPVATAVSFIDSKGTRVRMSASREI<br>LLCGGVYATPQLLMLSGVGPAEHLRSHGIEIVADVPAVGQNLQDHAAAMVSFESQNPEKDKANSSVYYTERTGKNIGTLLNYVFRGKGPLTSPMCEAGG<br>FAKTDPSMDACDLQLRFIPFVSEPDPYHSLADFATAGSYLQNRANRPTGFTIQSVAARPKSRGHVQLRSTDVRDSMSIHGNWISNDADLKTLVHGVKLC<br>RTIGNDDSMKEFRGRELYPGGEKVSDADIEAYIRDTCHTANAMVGTCRMGIGEQAAVDPALQVKGVARLRVVDSSVMPTLPGGQSGAPTMMIAEKG<br>ADLIRAAARQADAATVGAAA |
| **TrxA-GsFAP** | | *Galderia sulphuraria* | >TrxA-GsFAP<br>MGHHHHHHSDKIIHLTDDSFDTDVLKADGAILVDFWAEWCGPCKMIAPILDEIADEYQGKLTVAKLNIDQNPGTAPKYGIRGIPTLLLFKNGEVAATKVG<br>ALSKGQLKEFLDANLAGSGSGVSGVDLGTENLYFQSGFDRSREFDYVIVGGGAAGCVLASRLSEDKRSTVLLLEAGKEDENFYIHVPMGFPYLVGSDLDW<br>KYQSTSEERLLDRKIMWPRGKVLGGSHAISVMLYLRGSANDYREWERCGALGWGPEDVLPYYLKAENNLRGVGPYHGKGGPLTVSDLPSPNEMSHAFV<br>IAGQTLDIPYNKDFNDWSHSQEGIGLFQVTQANGKRVSPATAYLHPVRSRRNLFIETQRHVEKILFTKEIGSCPRAIGVSYINSNGKRERAMIRKEVVVSAG<br>AYGSPQLLMLSGIGPSNVLDSIGIPTVMPLEGVGKNLQDHFAVMFSCKSPDPSKDRKRRNLYYTDDTGKDWKTILTWFLSGKGPLTSTMCEAGAFLKTNP<br>IFNDPDLQLRFIPFFSEADPYFSLSDYSSKGMFFKNRSHRPSGFTIQSVAIRPKSRGRLTIQSKDPRMLPIIESGWFSSQEDLETLLRGISLSQKLVHASPLASYF<br>GEQCFPSTGLSKKEDIIRYISSTCHTANAVVGTCRMGTDKQAVVNPNLQVMGVERLRVLLQL |
| **TrxA-EsFAP** | | *Ectocarpus silicolusus* | >TrxA-EsFAP<br>MGHHHHHHSDKIIHLTDDSFDTDVLKADGAILVDFWAEWCGPCKMIAPILDEIADEYQGKLTVAKLNIDQNPGTAPKYGIRGIPTLLLFKNGEVAATKVG<br>ALSKGQLKEFLDANLAGSGSGVSGVDLGTENLYFQSSMSVAEEGHKFIIIGGGTAGCVLANRLSADKDNSVLVLEAGSEKFNDRNIKMPIAILRLFKSVFD<br>WGFQSENEKFATGDGIYLCRGKVLGGSSCTNVMLYHRGEEADYDAWGVDGWKGKDVLPYFKKAENNRSKKKGEFHGKGGLMQVENARYMNPLTKL<br>FFKACEQAGLSENEDFNDWSHSQEGFGRFQVAQKRGKRCSAASSYLKEAMGRKNLDVQTSAQITKVLIENGGAIGVEYVRDGEKKIAKLAVGGEILLAG<br>GAISSPQVLMLSGVGPAEHLRSKGIEVKSNVPGVGKNLRDHPAVTVMADINKPISITDKVLKEGSGDVNKITALQWLLTGTGPLTSPGCENGAFFKTTPDK<br>AAADLQLRFVPGRSTTPDGVKAYNTIGTKGRPPSGVTVQVVGIRPQSEGHVELRSSDPFDKPHIVTNYLESGEDMASLTNGIEMARKLFDQEAFGEMVD<br>KEVFPGRDNKEISEYIKSTVHSANALVGTCKMGEESDNMSVVNSALKVKGVAGLRVIDSSVMPSIPGGQTAAPTIMIAEKAADMLMA |
| **TrxA-NgFAP** | | *Nannochloropsis gaditana* | >TrxA-NgFAP<br>MGHHHHHHSDKIIHLTDDSFDTDVLKADGAILVDFWAEWCGPCKMIAPILDEIADEYQGKLTVAKLNIDQNPGTAPKYGIRGIPTLLLFKNGEVAATKVG<br>ALSKGQLKEFLDANLAGSGSGVSGVDLGTENLYFQSLQSVSMKAPAAVASSTYDYIIVGGGIGGCVLANRLTESGRFKVLLLEAGKSAERNPYVNIPAGVV<br>RLFKSALDWQFESAPERHLDGKEVYLVRGKAMGGSSAVNVMLVHRGSASDYAKWEAEGAQGWGPEEALRYFKKMEDNLVGGEGRWHGQGGMYP<br>VDDVKYQNPLSKRFLQACEEYGWRANPDFNDWSHPQDGYGSFKVAQKHGKRVTAASGYLNKAVRRRPNLDILSEALVTRVLLEGEGDVKAVGVEFTGK<br>DGKTHQVRTTGKAGEVLLAGGAVNSPQLLMLSGIGPEADLQAVGIATKVNRPGVGENLQDHPAVTIAHNITRPISLCDDLFLFHTPVPKPHQVLRWTLT<br>GSGPLTTPGCDHGAFLKTREDLQEPNVQFRFIAGRGSDPDGVRSYIMGGSARPLSGLTLQVVNIRPKSKGKLTLASKDPLKKPRIEVRYLSAAEDLQALRTG<br>MRIGRDLIKQRAFADILDEEVFPGPAAQTDEELDAYIRDSLHTANALVGTCKMGSVEDRNAVVDPECRVIGVGGLRVVDASVMPVIPGGQTGSGTTML<br>AEKAADLVRAHAGDLVEMGVQDEERKGGWFNGLLGRKQKVAT |
| **TrxA-AtTES** | Octanoyl-CoA specific thioesterase | *Anaerococcus tetradius* | >AtTES<br>MGSSHHHHHHSSGLVPRGSHMASMTGGQQMGRIRMKFKKKFKIGRMHVDPFNYISMRYLVALMNEVAFDQAEILEKDIDMKNLRWIIYSWDIQIEN<br>NIRLGEEIEITTIPTHMDKFYAYRDFIVESRGNILARAKATFLLMDITRLRPIKIPQNLSLAYGKENPIFDIYDMEIRNDLAFIRDIQLRRADLDNNFHINNAVYF<br>DLIKETVDIYDKDISYIKLIYRNEIRDKKQIQAFARREDKSIDFALRGEDGRDYCLGKIKTNV |
| **CpTES-287** | | *Cuphea palustris* | >CpTES-287<br>MGSSHHHHHHSSGLVPRGSHMASMTGGQQMGRIRMFDRKSKRPSMLMDSFGLERVVQDGLVFRQSFSIRSYEICADRTASMETVMNHVQETSLNQ<br>CKSIGLLDDGFGRSPEMCKRDLIWVVTRMKIMVNRYPTWGDTIEVSTWLSQSGKIGMGRDWLISDCNTGEILVRATSVYAMMNQKTRRFSKLPHEVR<br>QEFAPHFLDSPPAIEDNDGKLQKFDVKTGDSIRKGLTPGWYDLDVNQHVSNVKYIGWILESMPTEVLETQELCSLTLEYRRECGRDSVLESVTSMDPSKV<br>GDRFQYRHLLRLEDGADIMKGRTEWRPKNAGTNGAISTGKT |
| **ChTES** | | *Cuphea hookeriana* | >ChTES<br>MGSSHHHHHHSSGLVPRGSHMASMTGGQQMGRIRMVAAAASSAFFPVPAPGASPKPGKFGNWPSSLSPSFKPKSIPNGGFQVKANDSAHPKANGS<br>AVSLKSGSLNTQEDTSSSPPPRTFLHQLPDWSRLLTAITTVFVKSKRPDMHDRKSKRPDMLVDSFGLESTVQDGLVFRQSFSIRSYEIGTDRTASIETLMNH<br>LQETSLNHCKSTGILLDGFGRTLEMCKRDLIWVVIKMQIKVNRYPAWGDTVEINTRFSRLGKIGMGRDWLISDCNTGEILVRATSAYAMMNQKTRRLSK<br>LPYEVHQEIVPLFVDSPVIEDSDLKVHKFKVKTGDSIQKGLTPGWNDLDVNQHVSNVKYIGWILESMPTEVLETQELCSLALEYRRECGRDSVLESVTAMD<br>PSKVGVRSQYQHLLRLEDGTAIVNGATEWRPKNAGANGAISTGKTSNGNSVS |

**Table SA: Protein sequences of all the genes that were heterologous expressed in *E. coli* in this work**. The different FAP sequences were His tagged and cloned in *E. coli* without their respective chloroplast transit peptide and fuse or not to TrxA. **Purple:** His tag sequence, **Red:** TrxA sequence, Pink: TEV (tobacco etch virus protease recognition and cleavage site), **Green:** Linker aminoacids.

| Name of the plasmid | Features | Backbone | Size (bp) | References |
|---|---|---|---|---|
| pLIC03 (EV) | 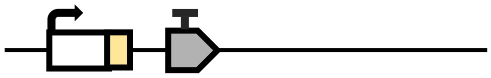 | pET28 | 7286 | Ali-Ahmad et al. 2021<br>Vincent et al. 2017 |
| pLIC03-CvFAP | 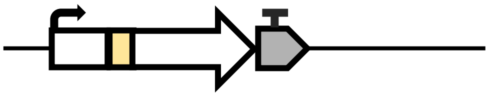 | pLIC03 | 7142 | This work |
| pLIC07 (EV) | 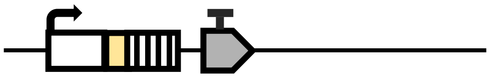 | pET28 | 7624 | Sorigué et al. 2017 |
| pLIC07-CvFAP | 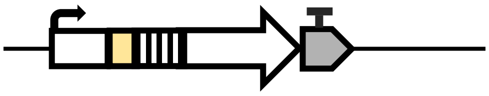 | pLIC07 | 7482 | Sorigué et al. 2017 |
| pLIC07-FAP homologs | 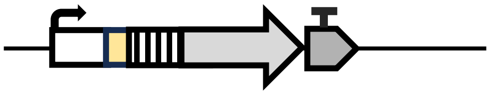 | pLIC07 | 7000-8000 | Moulin et al. 2021 |
| pDAWN (EV) | 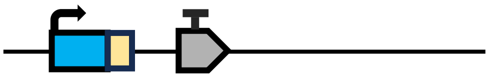 | pDAWN | 7213 | Oht al. 2012 |
| pDAWN-cvFAP | 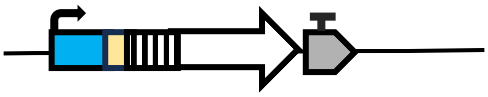 | pDAWN | 9371 | This work |
| pDAWN-C8_TES + cvFAP | 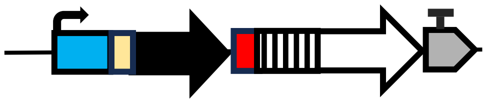 | pDAWN | 10081 - 1036 | This work |
| pDAWN-ChTES | 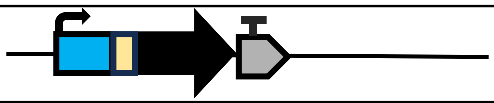 | pDAWN | 8438 | This work |

**Table SB:** **Genetic constructions used in this work in order express FAP and C8_TES genes in *E. coli* strains.** EV= Empty vector. All vectors provide resistance to kanamycin. Features: T7 promoter Histag CDS of Thioredoxin A Terminator CDS of cvFAP CDS of FAP homologs Blue-light inducible promoter CDS of C8_TES (AtTES, CpTES, ChTES) RBS (Ribosome binding site).

| Construction | Technique | Restriction enzyme | Backbone | Insert | Primer | Sequence | Observation |
|---|---|---|---|---|---|---|---|
| pLIC03-CvFAP | Golden-Gate | BsaI | pLIC03 | CvFAP gene | pLIC03-CvFAP fw | ttggtctcccaatggccagcgcagttgaagatattcg | - |
| | | | | | pLIC03-CvFAP rev | ttggtctcgtacttcatgctgcaacggttgccgg | |
| pLIC07-CvFAP | Inphusion | BsaI | pLIC07 | CvFAP gene | pLIC07-CvFAP fw | ctgtacttccaatcagccagcgcagttgaagatattc | - |
| | | | | | pLIC07-CvFAP rev | tatccacctttactgttatcatgctgcaacggttgccggtg | |
| pDAWN-CvFAP | Restriction digest and ligation | HindIII and XhoI | pDAWN | CvFAP gene fused with TrxA in 5' extremity | pDAWN-CvFAP fw | ttaagcttgagcgataaaattattcacctgactgacgac | - |
| | | | | | pDAWN-CvFAP rev | ttctcgagtcatgctgcaacggttgccggtg | |
| pDAWN-ChTES | Inphusion | NheI and NotI | pDAWN | ChTES gene | ChTES fw | cagccatatggctagcatgactggtggacagcaaatggg tcggatccgaatggttgcggccgcggccagc | Plasmid used for construction of other plasmids only. |
| | | | | | ChTES rev | ttaaagtacgcggccgcttagctaacgctattgccattgc tcgttttgcccgtgc | |
| pDAWN-ChTES + CvFAP | Restriction digest and ligation | NotI and XhoI | pDAWN-ChTES | CvFAP gene fused with TrxA and RBS in 5' extremity | ChTES + CvFAP fw | ttgcggccgcgtactttaactttaagaaggagatatac | - |
| | | | | | ChTES + CvFAP rev | ttctcgagtcatgctgcaacggttgccggtg | |
| pDAWN-CpTES-278 + CvFAP | Restriction digest and ligation | NheI and NotI | pDAWN-ChTES + CvFAP | CpTES-287 gene | - | - | Restriction sites are included in the CpTES-287 gene. |
| | | | | | - | - | |
| pDAWN-AtTES + CvFAP | Restriction digest and ligation | NheI and NotI | pDAWN-ChTES + CvFAP | AtTES gene | - | - | Restriction sites are included in the AtTES gene. |
| | | | | | - | - | |

**Table SC**: Molecular biology techniques and primer sequences used in this work. The AtTES and CpTES-287 genes were commanded with NheI and NotI sites in 5' and 3' ends respectively to facilitate cloning.

| *E. coli* strain name | Genetic background | Relevant genotype | Reference |
|---|---|---|---|
| BL21(DE3) | B834 | E. coli str. B $F^{-}$ ompT gal dcm lon hsd$S_B$($r_B^{-}m_B^{-}$) λ(DE3 [lacI lacUV5-T7p07 ind1 sam7 nin5]) [malB$^{+}$]$_{K-12}$($\lambda^{S}$) | – |
| Rosetta™(DE3) | BL21(DE3) | $F^{-}$ompT hsd$S_B$($r_B^{-}$ $m_B^{-}$) gal dcm (DE3) pRARE (Cam$^{R}$) | – |
| Rosetta-gami™ 2(DE3) | Rosetta™(DE3) | Δ(ara-leu)7697 ΔlacX74 ΔphoA PvuII phoR araD139 ahpC galE galK rpsL (DE3) F′[lac$^{+}$ lacI$^{q}$ pro] gor522::Tn10 trxB pRARE2 (Cam$^{R}$, Str$^{R}$, Tet$^{R}$ | – |
| C41(DE3) | BL21(DE3) | $F^{-}$ ompT gal dcm hsd$S_B$($r_B^{-}$ $m_B^{-}$)(DE3) | – |
| C43 (DE3) | C41(DE3) | F – ompT hsdSB (rB- mB-) gal dcm (DE3) | – |
| DH5α | K-12 | fhuA2 lac(del)U169 phoA glnV44 Φ80' lacZ(del)M15 gyrA96 recA1 relA1 endA1 thi-1 hsdR17 | – |
| BL21(DE3) pRIL | BL21(DE3) | pRIL in BL21(DE3), Cam$^{R}$ | – |
| BL21(DE3) pRARE2 | BL21(DE3) | pRARE2 in BL21(DE3), Cam$^{R}$ | – |
| DH5α pRIL | DH5α | pRIL in DH5α, Cam$^{R}$ | – |
| BL21(DE3) pRIL pLIC03 | BL21(DE3) pRIL | pLIC03 in BL21(DE3) pRIL, Cam$^{R}$ | – |
| BL21(DE3) pRIL pLIC03-CvFAP | | pLIC03-CvFAP in BL21(DE3) pRIL, Cam$^{R}$ | This work |
| BL21(DE3) pRIL pLIC07 | | pLIC07 in BL21(DE3) pRIL, Cam$^{R}$ | Sorigué *et al*. 2017 |
| BL21(DE3) pRIL pLIC07-CvFAP | | pLIC07-CvFAP in BL21(DE3) pRIL, Cam$^{R}$ | |
| BL21(DE3) pRIL pLIC07-CrFAP | | pLIC07-CrFAP in BL21(DE3) pRIL, | Moulin et al. 2021 |
| BL21(DE3) pRIL pLIC07-CcFAP | | pLIC07-CcFAP in BL21(DE3) pRIL | |
| BL21(DE3) pRIL pLIC07-GsFAP | | pLIC07-GsFAP in BL21(DE3) pRIL | |
| BL21(DE3) pRIL pLIC07-EsFAP | | pLIC07-EsFAP in BL21(DE3) pRIL | |
| BL21(DE3) pRIL pLIC07-NgFAP | | pLIC07-NgFAP in BL21(DE3) pRIL | |
| BL21(DE3) pRIL pDAWN | | pDAWN in BL21(DE3) pRIL | This work |
| BL21(DE3) pRIL pDAWN-CvFAP | | pDAWN-CvFAP in BL21(DE3) pRIL | |
| BL21(DE3) pRIL pDAWN-CvFAP+AtTES | | pDAWN-CvFAP+AtTES in BL21(DE3) pRIL | |
| BL21(DE3) pRIL pDAWN-CvFAP+CpTES | | pDAWN-CvFAP+CpTES in BL21(DE3) pRIL | |
| BL21(DE3) pRIL pDAWN-CvFAP+ChTES | | pDAWN-CvFAP+ChTES in BL21(DE3) pRIL | |
| BL21(DE3) pRIL pDAWN-ChTES | | pDAWN-ChTES in BL21(DE3) pRIL | |
| BL21(DE3) pRARE2 pDAWN-CvFAP+ChTES | | pDAWN-CvFAP+ChTES in BL21(DE3) pRIL | |
| BL21(DE3) pRARE2 pDAWN-CvFAP+ChTES | BL21(DE3) pRARE2 | pDAWN-CvFAP+ChTES in BL21(DE3) pRARE2 | |
| Rosetta(DE3)™ pDAWN-CvFAP+ChTES | Rosetta™(DE3) | pDAWN-CvFAP+ChTES in Rosetta™(DE3) | |
| Rosetta-gami™ 2(DE3) pDAWN-CvFAP+ChTES | Rosetta-gami™ 2(DE3) | pDAWN-CvFAP+ChTES in Rosetta-gami™ 2(DE3) | |
| C41(DE3) pRIL pDAWN-CvFAP+ChTES | C41(DE3) pRIL | pDAWN-CvFAP+ChTES in C41(DE3) pRIL | |
| C43(DE3) pRIL pDAWN-CvFAP+ChTES | C43(DE3) pRIL | pDAWN-CvFAP+ChTES in C43(DE3) pRIL | |
| DH5α pRIL pDAWN-CvFAP+ChTES | DH5α pRIL | pDAWN-CvFAP+ChTES in DH5α pRIL | |

**Table SD:** All *E. coli* strains used in this work.
